# Alternative Polyadenylation Determines the Functional Landscape of Inverted Alu Repeats

**DOI:** 10.1101/2023.11.30.569399

**Authors:** Jayoung Ku, Sujin Kim, Keonyong Lee, Doyeong Ku, Namwook Kim, Hyunsu Do, Hyeonjung Lee, Jinju Han, Young-suk Lee, Yoosik Kim

## Abstract

Inverted Alu repeats (IRAlus) are abundantly found in the transcriptome, especially in introns and 3′ UTRs. Yet, the biological significance of 3′ UTR IRAlus remains largely unknown. Here, we find that IRAlus induce the silencing of genes involved in essential signaling pathways. We utilize J2 antibody to directly capture and map the double-stranded RNA structure of 3′ UTR IRAlus in the transcriptome. Bioinformatic analysis reveals alternative polyadenylation as a major axis of IRAlus-mediated gene regulation. Notably, the expression of mouse double minute 2 (MDM2), an inhibitor of p53, is upregulated by the exclusion of IRAlus during UTR shortening, which is exploited to silence p53 during tumorigenesis. Moreover, the transcriptome-wide UTR lengthening in neural progenitor cells results in the global downregulation of genes associated with neurodegenerative diseases, such as amyotrophic lateral sclerosis, via IRAlus inclusion. Our study establishes the functional landscape of 3′ UTR IRAlus and its role in human pathophysiology.

## INTRODUCTION

Alu elements are primate-specific short interspersed nuclear elements (SINEs). With more than 1.1 million copies, Alus constitute greater than 10% of the human genome, making them the most abundant repetitive sequences.^1^ Interestingly, they are found primarily in gene-rich areas, like introns and untranslated regions (UTRs), rather than randomly distributed throughout the genome.^1^ Alu elements were originally considered as ‘selfish’ or junk DNA sequences due to their inert biological function except for their self-amplification through retrotransposition.^2,3^ However, when two Alus with opposite orientations are present in the vicinity, their high-sequence similarities allow them to form an intramolecular hairpin structure with a long double-stranded stem. Also known as inverse repeats of Alus (IRAlus), they constitute a major class of endogenous double-stranded RNAs (dsRNAs) that can activate innate immune response systems.^4–6^

IRAlus in the 3′ UTR adds an important layer of post-transcriptional gene regulation. They are recognized by dsRNA-dependent adenosine deaminase 1 (ADAR1), which catalyzes the deamination reaction on adenosine, converting it to inosine (A-to-I editing).^7^ These A-to-I hyper-edited RNAs, together with their long dsRNA structures, are bound by non-POU domain containing octamer binding (NONO) and splicing factor proline and glutamine-rich (SFPQ) proteins, which sequester the host mRNA with 3′ UTR IRAlus in nuclear paraspeckles.^8–10^ This nuclear sequestration reduces the translational efficiency of the mRNA, resulting in the gene silencing effect. These 3′ UTR IRAlus-containing mRNAs can escape the nuclear retention under certain conditions. For example, deficiency of nuclear-enriched abundant transcript 1 (NEAT1) long noncoding RNA (lncRNA) disrupts the paraspeckle assembly and facilitates the nuclear export of mRNAs with 3′ UTR IRAlus.^8^ In addition, STAUFEN 1 (STAU1) interacts with nuclear retained IRAlus elements and augments the nuclear export of host mRNAs.^11^ On the other hand, coactivator-associated arginine methyltransferase 1 (CARM1) regulates the nuclear retention by both suppressing *NEAT1* transcription and reducing the RNA binding ability of NONO by inducing protein methylation.^9^

Despite the clear gene silencing effect of 3′ UTR IRAlus, their biological function and pathophysiological relevance remain largely unknown. One of the reasons is that the list of genes with 3′ UTR IRAlus is still incomplete. Previous bioinformatics analysis identified 333 genes containing IRAlus in the 3′ UTR by analyzing human mRNA and expressed sequence tag (EST) sequences,^10^ but many of these mRNAs are not sequestered in the nucleus.^4^ In addition, most of the previous investigations were done using EGFP reporters. For instance, in human pituitary cells, both the nuclear retention of IRAlus reporter mRNA and the expression of the reporter protein follow a circadian rhythmic pattern due to the circadian expression pattern of *NEAT1* RNA and NONO protein.^12^ Yet, it is unclear whether genes involved in the circadian rhythm are regulated by 3′ UTR IRAlus.

In this study, we employed formaldehyde crosslinking immunoprecipitation and sequencing (fCLIP-seq) with a J2 antibody that recognized IRAlus dsRNA structure^5,13^ to directly capture and map mRNAs with functional 3′ UTR IRAlus at the transcriptome level. Our approach greatly expanded the number of genes regulated by IRAlus and improved the accuracy of the 3′ UTR IRAlus genes in terms of both the nuclear localization and translational suppression of the host mRNAs. Moreover, we downregulated a key 3′ end cleavage factor to induce global UTR lengthening and showed that IRAlus inclusion could regulate essential cell signaling pathways. Notably, such regulation by IRAlus played an important role during tumorigenesis that accompanied the transcriptome-wide 3′ UTR shortening event. We further analyzed IRAlus-mediated gene silencing in neural progenitor cells (NPCs) that naturally expressed long 3′ UTR variants to study the cell-specific gene regulation by IRAlus. Collectively, our work provides a comprehensive transcriptome analysis of the crosstalk between APA and 3′ UTR IRAlus and suggests IRAlus-mediated gene silencing as a key post-transcriptional regulatory mechanism exploited in human pathophysiology.

## RESULTS

### Identification of functional 3′ UTR IRAlus genes

The gene silencing effect of 3′ UTR IRAlus is mediated by interaction with nuclear paraspeckle proteins and *NEAT1* lncRNA, which sequester the host mRNA in the nucleus (Figure 1A).^8,10^ Indeed, when the subcellular localization of IRAlus-containing mRNAs (see method for the identification of 3′ UTR IRAlus genes) was examined using RNA-fluorescent *in situ* hybridization (RNA-FISH), *ZNF587* and *MCM4* mRNAs showed strong nuclear signal (Figure 1B). However, some mRNAs with 3′ UTR IRAlus, such as *RPL13* and *RPL15* mRNAs, exhibited primarily cytosolic expression (Figure 1B). Of note, the nuclear paraspeckle was visualized through *NEAT1* lncRNA, while *GAPDH* mRNA was used as a control for an mRNA without IRAlus (noIRAlus) that was localized in the cytosol (Figure 1B).

**Figure 1.**
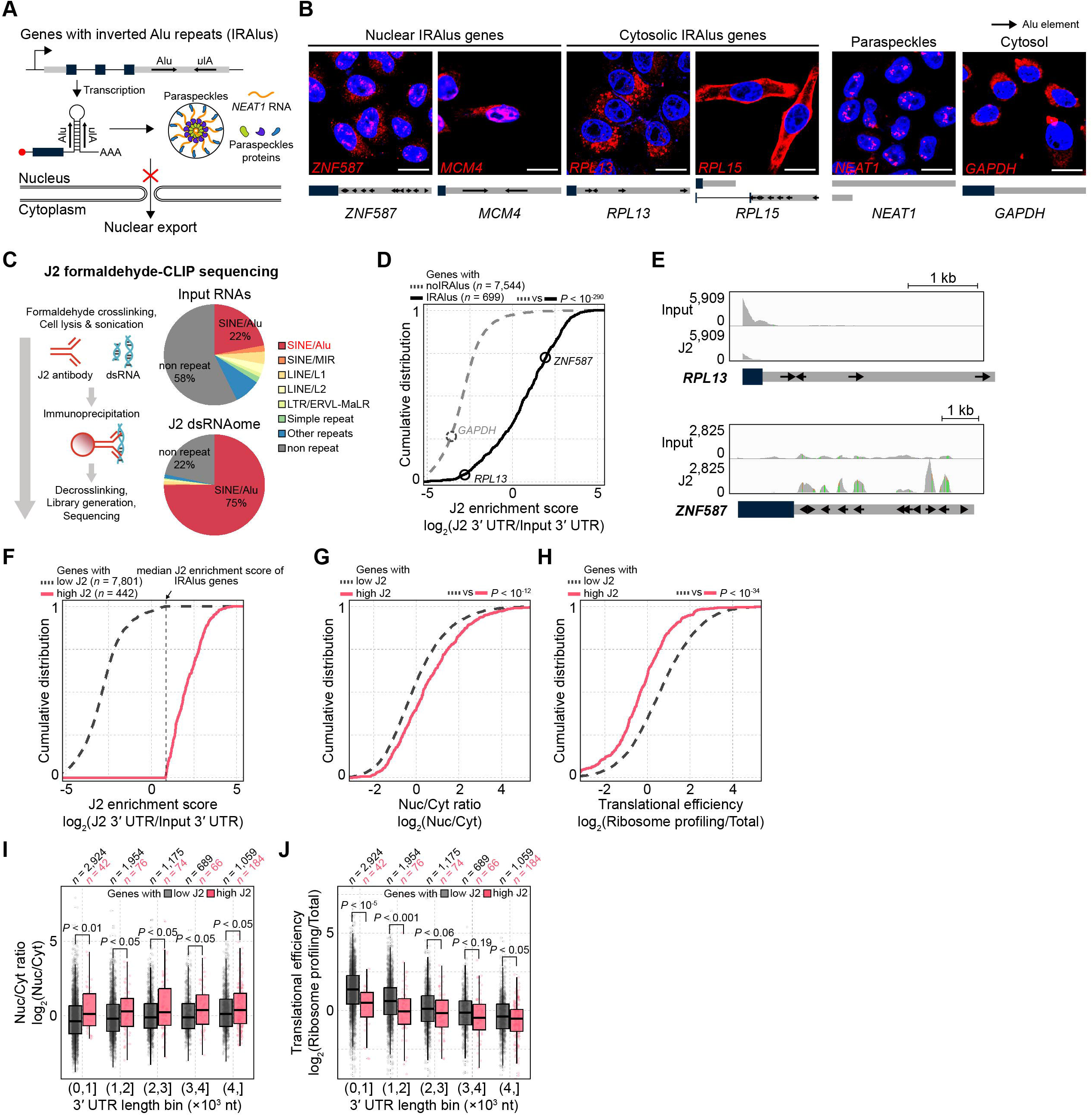
Identification of functional 3′ UTR IRAlus genes. (A) A schematic for the 3′ UTR IRAlus-mediated gene silencing. (B) RNA-FISH analysis for the subcellular localization of mRNAs of genes with 3′ UTR IRAlus in HeLa cells. The bar indicates 20 μm. The gray box indicates the 3′ UTR with arrows representing Alus. (C) Experimental scheme of J2 fCLIP-seq and the relative proportion of J2 fCLIP-seq reads by repetitive elements. (D) Cumulative distribution of the J2 enrichment scores of 3′ UTR IRAlus. (E) Representative example of 3′ UTR IRAlus genes showing different subcellular localization (*RPL13*, cytosol; *ZNF587*, nucleus). IGV snapshot of J2 fCLIP-seq reads with Alu element annotation of the 3′ UTR are shown. (F) Subclassification of genes based on the J2 scores. For the threshold, the median J2 score of IRAlus genes was used. (G and H) Cumulative distribution of the Nuc/Cyt ratio (G) and the translational efficiency (H) of mRNAs with the indicated J2 scores. (I and J) Comparison of the Nuc/Cyt ratio (I) and the translational efficiency (J) of mRNAs with the indicated J2 scores by matched 3′ UTR length bins. The *P* was calculated using the two-sided Mann-Whitney test. Boxplots show the 25th, 50th, and 75th percentiles. Whiskers represent a 1.5× interquartile range.

We employed J2 antibody that could recognize IRAlus elements to determine a list of genes with “functional” IRAlus elements in the 3′ UTR. First, we reanalyzed J2 fCLIP-seq data performed in HeLa cells that yielded about 75% of the sequencing reads mapped to SINE/Alu, most of which were IRAlus (Figure 1C). We then devised a J2 enrichment scoring strategy by taking the ratio of read counts of J2 fCLIP-seq in the 3′ UTR to those of input RNAs of the immunoprecipitation. Consistent with the idea that J2 could efficiently capture the IRAlus element, our analysis revealed significantly higher J2 enrichment scores for IRAlus genes compared to those of noIRAlus genes (Figure 1D). Notably, *RPL13* showed a low J2 enrichment score while *ZNF587* showed a high J2 enrichment score. When the sequencing reads accumulation at *RPL13* and *ZNF587* loci were visualized through IGV, it was clear that *RPL13* mRNA in HeLa cells did not express a long 3′ UTR variant containing the IRAlus while *ZNF587* mRNA did express a UTR variant with IRAlus elements (Figure 1E). Therefore, analysis of J2 fCLIP-seq data could generate a more accurate IRAlus gene list by incorporating the information on UTR variants abundantly expressed in the cell of interest.

Next, we used the median J2 enrichment score for the annotation-based IRAlus genes as a threshold to determine the functional IRAlus genes with high J2 enrichment scores, which yielded a total of 442 genes (Figure 1F). To analyze their nuclear retention, we performed mRNA-seq following subcellular fractionation to isolate nuclear and free cytosolic fractions in HeLa cells (Figure S1A). We then calculated the nuclear/cytosol (Nuc/Cyt) ratio of mRNAs with high J2 scores, which yielded significant *P* in both Nuc/Cyt ratio and the translational efficiency (Figures 1G and 1H). Of note, translational efficiency was determined by reanalyzing published ribosome profiling data in HeLa cells.^14^ In addition, the J2 enrichment score correlated with the Nuc/Cyt ratio (Spearman *r* = 0.11, *P* < 10^-12^; Figure S1B) and the translational repression (Spearman *r* = -0.20, *P* < 10^-43^; Figure S1C) of mRNAs.

In addition to hosting IRAlus elements, 3′ UTR provides a platform for diverse post-transcriptional gene regulations, including RNA interference by microRNAs (miRNAs).^15^ To rule out the possibility that the stronger gene silencing effect of high J2 genes was attributable to increased interactions with miRNAs, we compared 1) the probability of conserved targeting (Aggregate P_CT_ ^16^) and 2) the expected repression (CWCS^17^) of abundant miRNAs in HeLa cells (>1% abundance of total miRNAs, *n* = 22; Figure S1D).^18^ We found that the gene silencing effect of high J2 genes was independent of miRNA binding (Figures S1E and S1F). In addition, we analyzed the potential effect of UTR length. Indeed, high J2 genes exhibited a longer 3′ UTR length distribution (Figure S1G) and less RNA stability (Figure S1H). However, when we compared low and high J2 genes after binning them with a specific range of 3′ UTR length, we still found a significant gene silencing effect in high J2 genes, indicating that IRAlus are major regulatory factors for the nuclear retention of mRNAs (Figures 1I and 1J).

### Classification of the 3′ UTR IRAlus genes

Our analysis revealed that out of 699 IRAlus-containing genes abundantly expressed in HeLa cells, 422 of them showed high J2 enrichment scores. One possible explanation is that certain genes may express a short 3′ UTR variant without IRAlus in HeLa cells due to alternative polyadenylation (APA). APA in the 3′ UTR produces largely two mRNA variants with different lengths and sequences. The 3′ UTR upstream of the proximal poly(A) site is generally named the common UTR (cUTR), and the rest of the 3′ UTR is named the alternative UTR (aUTR).^15^ To better understand the biological significance of IRAlus-mediated gene silencing, it is important to incorporate the potential effect of APA. Therefore, we subclassified these 422 high J2 genes based on their cell-specific mode of regulation based on the APA information.

We utilized the strong poly(A) cleavage site information from the PolyASite 2.0^19^ to determine whether the IRAlus elements were present in major or minor 3′ UTR variants (Figure 2A). If a gene contains an IRAlus element in the major 3′ UTR variant defined by the strong poly(A) site, then we consider it a “constitutive IRAlus gene” (Figure 2B, Case #1, *n* = 191). If a gene contains at least one Alu for the IRAlus pair in a minor 3′ UTR variant, then it is considered a “conditional IRAlus gene” (Figure 2B, Case #2, *n* = 158). Of note, these conditional IRAlus genes with high J2 enrichment scores express IRAlus-containing 3′ UTR variants in HeLa cells and are likely to induce cell-specific IRAlus-mediated gene regulation. For example, *ZNF791* was predicted to contain IRAlus elements in the aUTR (minor 3′ UTR variant) (Figure 2C). However, raw J2 fCLIP-seq reads showed the predominant usage of distal poly(A) site with strong J2 enrichment of its aUTR in HeLa cells (Figure 2C). In addition, when a gene contains an IRAlus element within the 2,000 nt beyond the annotated 3′ end before reaching the neighboring gene, we categorize it as an “extended IRAlus gene” (Figure 2B, Case #3, *n* = 64). Of note, 2,000 mer was chosen as the limit because it reflects twice the median 3′ UTR length of all analyzed genes. An example of the extended IRAlus gene is *KRBA2* which expressed a longer 3′ UTR than the RefSeq annotation in HeLa cells, with strong J2 enrichment at Alus in this region (Figure 2D).

**Figure 2.**
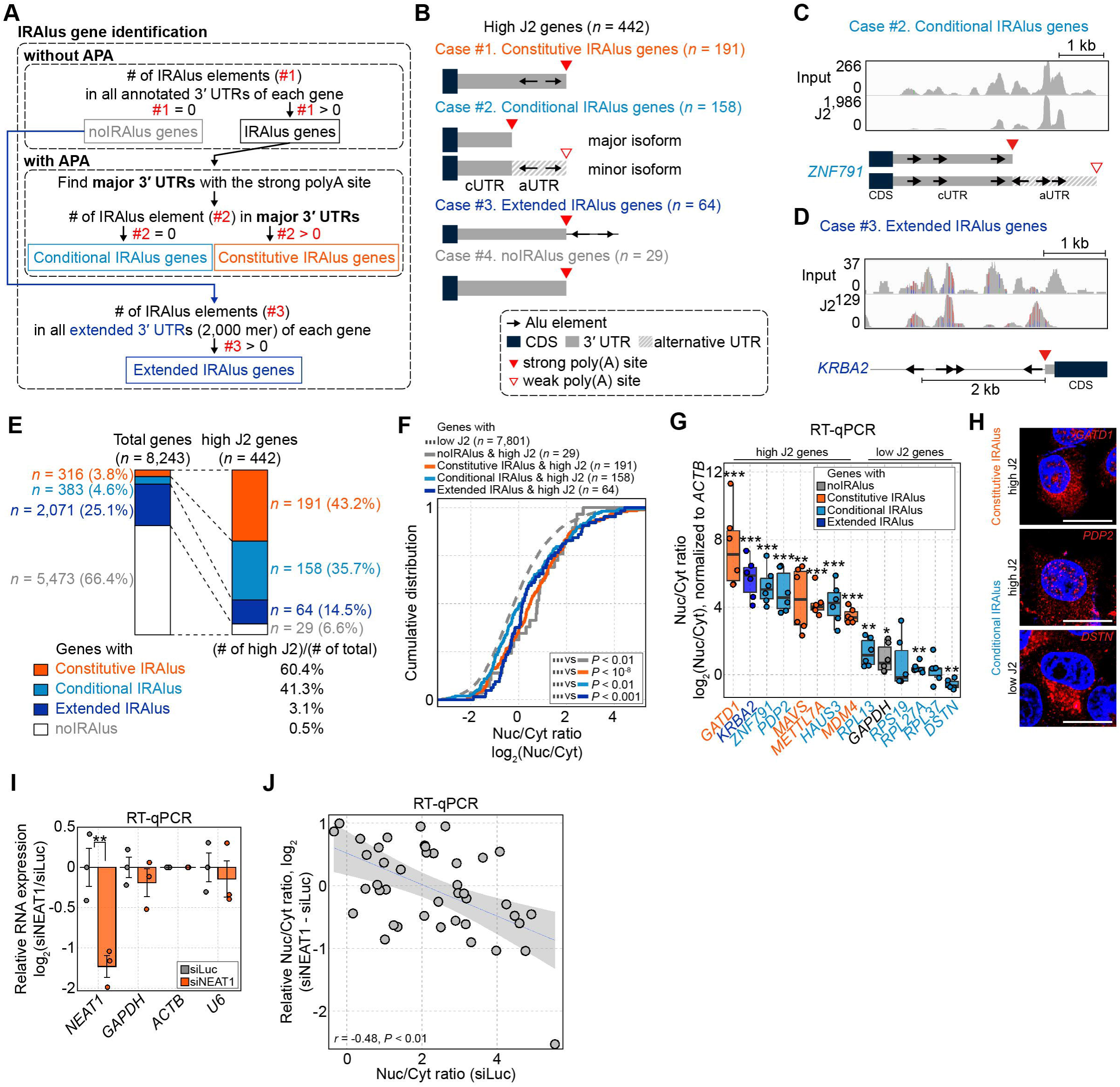
Classification and validation of 3′ UTR IRAlus genes. (A) Overview of the classification of 3′ UTR IRAlus genes using APA information. (B) Four categories of high J2 genes based on the location of IRAlus. (C and D) Representative example of conditional (*ZNF791*; C) and extended (*KRBA2*; D) IRAlus genes with high J2 scores. IGV snapshots of J2 fCLIP-seq reads mapped to the 3′ UTR are shown. (E) Distribution of the IRAlus genes in total (*n* = 8,243) or high J2 (*n* = 442) genes. The relative percentage of each IRAlus gene category in high J2 genes compared to the total genes is shown. (F) Cumulative distribution of the Nuc/Cyt ratio of mRNAs with the indicated 3′ UTR IRAlus and J2 scores. The *P* was calculated using the two-sided Mann-Whitney test. (G) The Nuc/Cyt ratio of the representative mRNAs with the indicated 3′ UTR IRAlus and J2 scores analyzed by RT-qPCR (*n* = 6). The *P* was calculated using the one-sided Student’s t-test. (H) RNA-FISH analysis of *GATD1*, *PDP2*, and *DSTN* mRNAs. The bar indicates 20 μm. (I) Knockdown efficiency analyzed by RT-qPCR in HeLa cells transfected with siNEAT1 for 2 days (*n* = 3). Data are represented as mean ± s.e.m. The *P* was calculated using the one-sided Student’s t-test. (J) Scatter plot for comparison between basal Nuc/Cyt ratio (x-axis) and relative Nuc/Cyt ratio (y-axis) of mRNAs of genes from (G) analyzed by RT-qPCR in cells transfected with siNEAT1 for 2 days (*n* = 3). Correlation *r* and its *P* was calculated using the Spearman correlation. Boxplots show the 25th, 50th, and 75th percentiles. Whiskers represent a 1.5× interquartile range. \**P* < 0.05, \*\**P* < 0.01, \*\*\**P* < 0.001.

Interestingly, some of the high J2 genes (29 out of 442) did not contain IRAlus elements in their annotated or extended 3′ UTRs (Figure 2B, Case #4, *n* = 29). When examining these genes in detail, most of them exhibited a complex splicing pattern that is hard to distinguish introns and 3′ UTRs. In addition, the *NBPF3* gene, one of the noIRAlus genes, included a dsRNA structure consisting of a long interspersed nuclear element (LINE/L1) (Figure S2A). Another notable noIRAlus showed an SVA element in the vicinity of an Alu element. SVA elements are one member of the non-LTR (long terminal repeat) retrotransposon family of 0.7-4 kb in length.^20^ Because they include an antisense Alu-like sequence (Figure S2B), Alu and SVA with the same orientation can form IRAlus-like dsRNA structure. For example, *TRIM56* 3′ UTR contained an SVA_C retrotransposon near an AluSq2 element with the same orientation, which resulted in strong J2 sequencing read accumulation (Figure S2C). The long dsRNA structure predicted by RNAfold near the sequencing peak further supports the potential Alu-SVA interaction to generate a long dsRNA structure (Figure S2C). Interestingly, *TRIM56* mRNA also showed strong nuclear retention like a typical 3′ UTR IRAlus gene (Figure S2D).

When we apply the same classification category to 8,243 abundantly expressed genes in HeLa cells, we find 316 constitutive IRAlus, 383 conditional IRAlus, 2,071 extended IRAlus, and 5,473 noIRAlus genes (Figure 2E). Out of 316 annotated constitutive IRAlus, 60.4% of genes show high J2 enrichment scores. For conditional and extended cases, high J2 genes only occupy 41.3% and 3.1%, respectively. This discrepancy is mostly from the PolyASite 2.0 database lacking HeLa cell-specific poly(A) cleavage information. Our analysis indicates that a large number of IRAlus-containing genes are likely to be regulated in a tissue-specific manner through APA and have the potential to be regulated by IRAlus in cells other than HeLa.

With this classification, we next asked whether a high J2 enrichment score could capture the functionality of the IRAlus element. In all IRAlus gene categories, genes with high J2 enrichment scores showed significantly higher Nuc/Cyt ratios (Figure 2F). Of note, if we did not use J2 enrichment scores, Nuc/Cyt ratios of conditional and extended IRAlus genes were indistinguishable from that of noIRAlus, suggesting that high J2 enrichment score specifically identify the functional 3′ UTR IRAlus (Figure S2E). We further confirmed the strong nuclear retention of mRNAs of constitutive IRAlus genes as well as conditional IRAlus genes with high J2 enrichment scores in HeLa cells using RT-qPCR following subcellular fractionation (Figure 2G) and RNA-FISH experiments (Figure 2H). To validate that mRNAs of IRAlus genes are sequestered in paraspeckles, we knockdowned *NEAT1* lncRNA and found that IRAlus-containing mRNAs were released from the nucleus (Figures 2I and 2J). Moreover, the negative correlation between the relative Nuc/Cyt ratio change in *NEAT1*-deficient cells and their basal Nuc/Cyt ratio (*r* = -0.48, *P* < 0.01) confirmed that strong nuclear retention of mRNAs of IRAlus genes was associated with *NEAT1* lncRNA (Figure 2J). Collectively, our analysis revealed that J2 fCLIP-seq analysis successfully identified HeLa-specific conditional and extended IRAlus genes with strong nuclear sequestration as well as LINE and Alu-SVA interaction that generated IRAlus-like functional element in the 3′ UTR.

### Regulation of 3′ UTR IRAlus-mediated gene silencing via UTR lengthening

Considering that most of the genes with high J2 enrichment scores are conditional and extended IRAlus, we set out to investigate the biological significance of these regulated IRAlus genes by modulating the 3′ UTR length. During tumorigenesis, transcriptome-wide 3′ UTR shortening occurs due to the increased expression of cleavage stimulation subunit 2 (CSTF2), a key 3′ end cleavage factor.^21^ Previous studies demonstrated that the co-depletion of *CSTF2* and its paralog *CSTF2T* resulted in global 3′ UTR lengthening by increased usage of distal poly(A) sites.^21^ Therefore, we utilized the double knockdown of *CSTF2* and *CSTF2T* to examine the downstream effect of IRAlus inclusion during UTR lengthening.

We performed both J2 fCLIP-seq and mRNA-seq on cytosolic and nuclear fractions in A549 cells transfected with a mixture of siCSTF2 and siCSTF2T. The knockdown efficiency was confirmed using RT-qPCR (Figure S3A). Of note, we chose A549 cells as a model cell line because the upregulation of *CSTF2* expression and its correlation with global 3′ UTR shortening events have been reported in lung cancer.^22^ As expected, the knockdown of *CSTF2* and *CSTF2T* resulted in a significant increase in J2 enrichment scores, regardless of the type of 3′ UTR IRAlus gene, resulting in an increased number of genes with high J2 enrichment scores (Figures 3A and S3B). Similar to HeLa cells, the J2 enrichment score successfully captured functional IRAlus elements in A549 cells (Figure 3B). We further found that the mRNAs of genes with constitutive or conditional IRAlus showed stronger nuclear retention in the depletion of *CSTF2* and *CSTF2T*, which was further confirmed using RT-qPCR following subcellular fractionation (Figures 3C and 3D).

**Figure 3.**
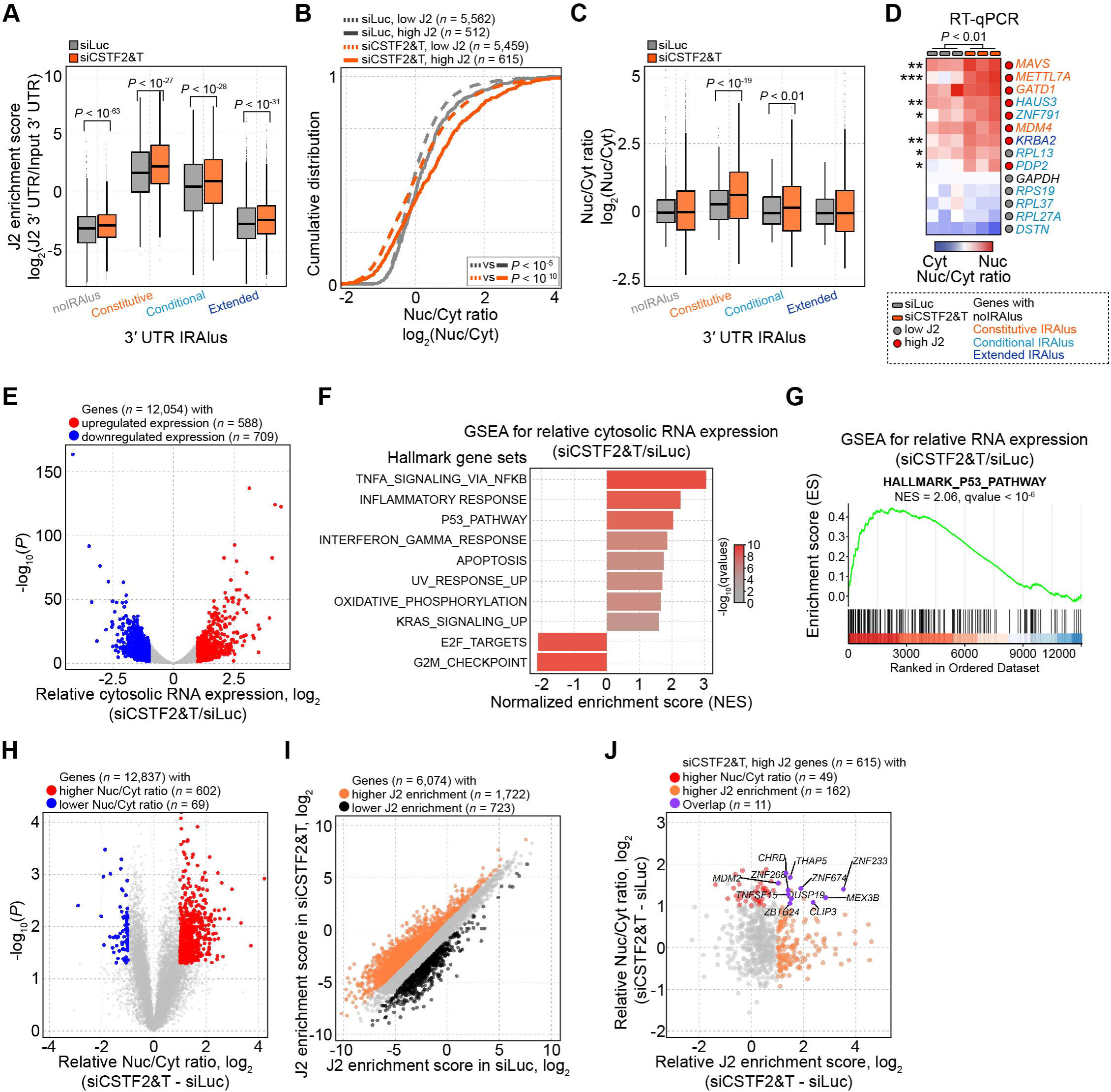
Regulation of IRAlus-mediated gene silencing via UTR lengthening. (A) J2 scores of indicated 3′ UTR IRAlus genes in A549 cells transfected with a mixture of siCSTF2 and siCSTF2T for 4 days. The *P* was calculated using the two-sided Wilcoxon signed rank test. (B) Cumulative distribution of the Nuc/Cyt ratio of mRNAs with the indicated J2 scores. The *P* was calculated using the two-sided Mann-Whitney test. (C) The Nuc/Cyt ratio of mRNAs with the indicated 3′ UTR IRAlus in A549 cells transfected with a mixture of siCSTF2 and siCSTF2T. The *P* was calculated using the two-sided Wilcoxon signed rank test. (D) The Nuc/Cyt ratio of selected mRNAs analyzed by RT-qPCR (*n* = 3). The *P* was calculated using the two-sided unpaired Student’s t-test. \**P* < 0.05 \*\**P* < 0.01 \*\*\**P* < 0.001. (E) Volcano plot of relative cytosolic RNA expression in A549 cells transfected with a mixture of siCSTF2 and siCSTF2T. The *P* was calculated using DESeq2 software. Significant cytosolic RNA expression change was defined as a |log_2_ fold change of cytosolic RNA expression| > 1 and *P* < 0.05, and indicated with colored dots (*n* = 1,297). (F and G) Significantly enriched Hallmark gene sets identified by GSEA for relative gene expression in A549 cells transfected with a mixture of siCSTF2 and siCSTF2T (F) and the GSEA results for the HALLMARK_P53_PATHWAY gene set (G). (H) Volcano plot of relative Nuc/Cyt ratio of mRNAs in CSTF2- and CSTF2T-deficient A549 cells. The *P* was calculated using the two-sided unpaired Student’s t-test. Significant Nuc/Cyt ratio change was defined as a |log_2_ fold change of Nuc/Cyt ratio| > 1 and *P* < 0.05, and indicated with colored dots (*n* = 671). (I) Scatter plot for J2 scores of mRNAs in A549 cells transfected with a mixture of siCSTF2 and siCSTF2T. Significant J2 enrichment score change was defined as a |log_2_ fold change of J2 enrichment score| > 1 and indicated with colored dots (*n* = 2,445). (J) Scatter plot for relative J2 enrichment score (x-axis) and relative Nuc/Cyt ratio (y-axis) of mRNAs.

We then performed gene-set enrichment analysis (GSEA) with pre-defined Hallmark gene sets^23^ using relative cytosolic mRNA abundance in A549 cells transfected with a siRNA mixture against *CSTF2* and *CSTF2T* (Figures 3E and 3F). The majority of activated pathways were related to inflammatory responses, such as tumor necrosis factor-alpha (TNF-α) signaling via nuclear factor kappa B (NF-κB). NF-κB is a family of transcription factors associated with immune and inflammatory responses.^24^ One of the upstream activators of NF-κB is protein kinase R (PKR),^25^ which can be activated by directly binding to IRAlus in 3′ UTRs.^4,11^ Increased level of phosphorylated PKR and its downstream eukaryotic initiation factor 2 alpha (eIF2α) upon *CSTF2* and *CSTF2T* knockdown confirmed that accumulated IRAlus element by 3′ UTR lengthening likely activated NF-κB pathway via PKR (Figure S3C). In addition to the NF-κB activation, GSEA analysis revealed responses to increased levels of endogenous dsRNAs, such as interferon-gamma response and apoptosis (Figure 3F).

To our surprise, genes involved in the downstream of p53 signaling pathway activation were enriched during global 3′ UTR lengthening (Figures 3F, 3G, and S3D). p53 is a well-known tumor suppressor protein and a transcription factor that controls cell cycle, DNA damage response, and apoptosis.^26^ To investigate whether the activation of the p53 pathway was related to altered IRAlus-mediated gene silencing effects, we examined genes with high J2 enrichment scores and strong nuclear sequestration in response to the knockdown of *CSTF2* and *CSTF2T* (Figures 3H-J). Among these genes, we found that mouse double minute 2 (*MDM2)*, an inhibitor of p53, exhibited a strong gene silencing effect in *CSTF2*- and *CSTF2T*-deficient A549 cells.

### Activation of the p53 pathway via silencing of MDM2

MDM2 is an E3 ubiquitin-protein ligase and maintains a low level of p53 through ubiquitin-mediated proteolysis of the protein by the 26S proteasome.^27^ By inhibiting p53, MDM2 acts as an oncogene, and its expression is upregulated in diverse types of cancer with wild-type *TP53*.^28,29^ We asked whether IRAlus inclusion could silence MDM2 expression, thus activating the p53 signaling. First, we analyzed previously published iCLIP data^21^ and found a putative CSTF2 binding region at the 3′ end of the cUTR of *MDM2* mRNA, indicating that CSTF2 could determine the 3′ end of the *MDM2* mRNA (Figure 4A). In addition, the aUTR of *MDM2* mRNA contained a clustered IRAlus-rich region (IRR) that showed significant J2 enrichment upon knockdown of *CSTF2* and *CSTF2T* (Figure 4B). When we analyzed the nuclear and cytosolic mRNA-seq data, we found increased nuclear retention of *MDM2* mRNA in cells deficient in *CSTF2* and *CSTF2T* (Figure 4C). Notably, the aUTR region with IRR showed even stronger sequencing read accumulation in the nuclear fraction, consistent with the long UTR (aUTR) variant of *MDM2* with IRR being sequestered in the nucleus (Figure 4C).

**Figure 4.**
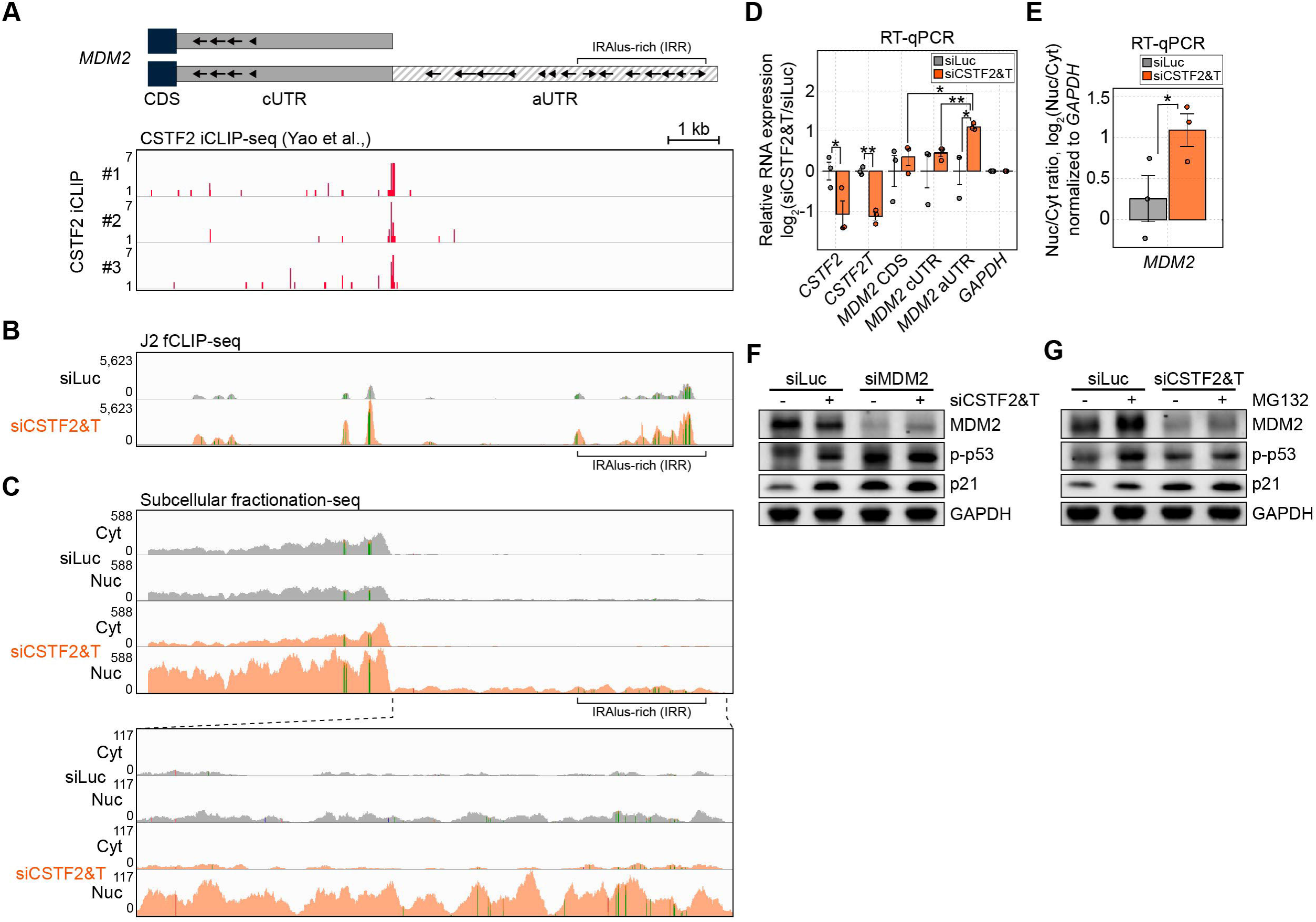
Suppression of the p53 pathway by MDM2 during UTR lengthening. (A-C) IGV snapshot of RNA-seq reads of the *MDM2* locus from CSTF2 iCLIP-seq (A), J2 fCLIP-seq (B), and nuclear and cytosolic mRNA-seq (C). Alu elements are indicated with arrows. (D) Knockdown efficiency and relative RNA expression of different genomic positions of MDM2 (CDS, cUTR, and aUTR) analyzed by RT-qPCR in A549 cells transfected with a mixture of siCSTF2 and siCSTF2T (*n* = 3). Data are represented as mean ± s.e.m. (E) The Nuc/Cyt ratio of *MDM2* mRNA analyzed by RT-qPCR in CSTF2- and CSTF2-deficient A549 cells (*n* = 3). Data are represented as mean ± s.e.m. (F) The effect of CSTF2-, CSTF2T-, and/or MDM2-deficiency on the activation of the p53 pathway. (G) The effect of MG132 treatment on activation of the p53 pathway in A549 cells transfected with a mixture of siCSTF2 and siCSTF2T. MG132 was treated one day before the cell harvest. The *P* was calculated using the one-sided Student’s t-test with unequal variance. \**P* < 0.05, \*\**P* < 0.01, \*\*\**P* < 0.001.

Next, we validated our mRNA-seq analysis of *MDM2* APA through RT-qPCR using primers targeting the coding sequence (CDS), cUTR, or aUTR of *MDM2* (Figure 4D). Increased expression of *MDM2* aUTR relative to that of *MDM2* CDS or cUTR confirmed the 3′ UTR lengthening of *MDM2* by *CSTF2* and *CSTF2T* knockdown. We also confirmed increased nuclear retention of *MDM2* mRNA when the 3′ UTR was lengthened (Figure 4E). Moreover, we observed the suppression of MDM2 protein expression and subsequent activation of the p53 pathway (Figure 4F). Of note, we knocked down MDM2 using siRNAs to confirm that the decreased expression of MDM2 resulted in the activation of the p53 pathway in A549 cells without external stimuli (Figure 4F). More importantly, when MDM2 expression was depleted, *CSTF2* and *CSTF2T* knockdown did not activate p53, suggesting that the regulation of p53 by *CSTF2* and *CSTF2T* deficiency was mediated by MDM2 (Figure 4F). Similarly, we examined the effect of inhibiting proteasomal degradation on *CSTF2/CSTF2T-*mediated p53 activation. We treated cells with MG132 to inhibit proteasomal degradation, which resulted in the activation of the p53 pathway despite the accumulation of MDM2 protein (Figure 4G). Of note, increased MDM2 expression by MG132 treatment is consistent with the previous studies showing that MDM2 is ubiquitinated by other ubiquitin ligases or autoubiquitination.^30^ We found that MG132 treatment did not enhance the p53 activation when *CSTF2* and *CSTF2T* were depleted, indicating that proteasomal activity was required for the activation of p53 in *CSTF2* and *CSTF2T* depleted cells (Figure 4G).

### Regulation of p53 signaling by IRAlus of *MDM2* 3′ UTR

To further investigate the importance of the IRAlus elements of the *MDM2* aUTR for the nuclear retention and subsequent silencing of MDM2, we employed the CRISPR system to remove IRR from the *MDM2* locus and analyzed its downstream effect. Using a total of four sgRNAs, two each targeting upstream and downstream of the IRR, we successfully generated a heterozygous A549 ΔIRR cell line (Figures 5A and S4A). Although a clone with a homozygous deletion was generated at the initial stage of genomic deletion (Figure S4B), it was not viable. We first confirmed that the knockdown of *CSTF2* and *CSTF2T* resulted in 3′ UTR lengthening of *MDM2* mRNA in both A549 WT and A549 ΔIRR cells using multiplex PCR (Figure S4C). Interestingly, the increased nuclear retention of *MDM2* mRNA by UTR lengthening was attenuated in A549 ΔIRR cells (Figures 5B, 5C, and S4D). Moreover, the activation of the p53 pathway by 3′ UTR lengthening in terms of both phosphorylated p53 expression and the induction of p53 downstream targets was also reduced in A549 ΔIRR cells (Figures 5D and 5E), which resulted in increased cell viability (Figure S4E). These results indicate the importance of the IRAlus elements of the *MDM2* aUTR in the p53 activation during 3′ UTR lengthening.

**Figure 5.**
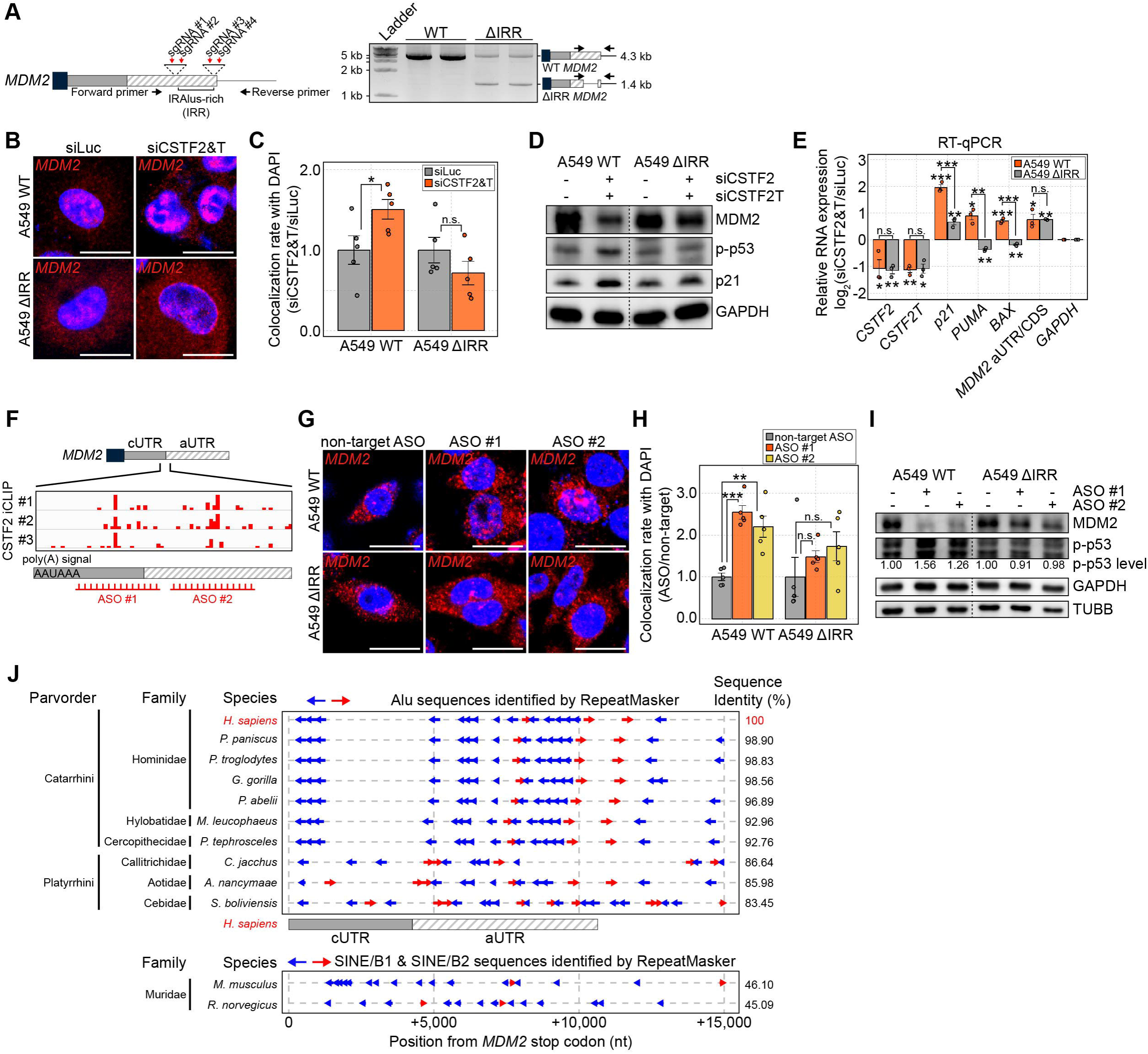
Regulation of p53 signaling by the *MDM2* 3′ UTR Alu cluster. (A) Schematic of Cas9 mediated genomic deletion of IRR of MDM2 (left panel) and PCR amplicons using gDNA extracted from A549 WT or A549 ΔIRR cells analyzed on a 1% TAE agarose gel (right panel). The expected migration of PCR amplicons is shown on the right along with their size. (B and C) RNA-FISH analysis of *MDM2* mRNA in A549 WT or A549 ΔIRR cells transfected with a mixture of siCSTF2 and siCSTF2T (B) and colocalization rate with DAPI (C; *n* = 5). The bar indicates 20 μm. (D and E) Repression of MDM2 expression and activation of the p53 pathway analyzed by western blotting (D) and RT-qPCR (E; *n* = 3) in A549 WT or A549 ΔIRR cells. (F) Schematic drawing of ASO design to target the CSTF2 binding sites of *MDM2* mRNA. (G and H) RNA-FISH analysis of *MDM2* mRNA in A549 WT or A549 ΔIRR cells transfected with an ASO targeting CSTF2 binding sites of *MDM2* mRNA for 2 days (G) and colocalization rate with DAPI (H; *n* = 5). The bar indicates 20 μm. (I) Repression of MDM2 expression and activation of the p53 pathway analyzed in A549 WT or A549 ΔIRR cells transfected with an ASO targeting CSTF2 binding sites of *MDM2* mRNA. (J) Alu or Alu-like element identified by RepeatMasker from the *MDM2* 3′ UTR of primates and murine. The sequences of 15,000 bp from the stop codon of *MDM2* were analyzed. For murine, SINE/B1 and SINE/B2 elements were used instead of Alus. The *P* was calculated using the one-sided Student’s t-test with unequal variance. n.s. not significant, \**P* < 0.05, \*\**P* < 0.01, \*\*\**P* < 0.001. All data are represented as mean ± s.e.m.

Next, we employed antisense oligonucleotides (ASOs) to specifically induce UTR lengthening of *MDM2* mRNA and examined its effect on MDM2 expression and p53 activation. We designed 11 ASOs to target the CSTF2 binding site of *MDM2*. All ASOs were 25-nucleotide in length with a phosphorothioate backbone and 2′-OMethyl sugar ring. We transfected these ASOs in A549 WT cells and identified two that showed a strong effect on 3′ UTR lengthening of *MDM2* (Figures 5F and S4F). Of note, we used non-target ASO,^31^ which did not induce 3′ UTR lengthening of *MDM2* mRNA, as a negative control. We found that both ASOs significantly increased the nuclear retention of *MDM2* mRNA (Figures 5G and 5H) and activation of the p53 pathway (Figure 5I) in A549 WT cells. However, these ASOs failed to increase *MDM2* mRNA nuclear retention and activate p53 signaling in A549 ΔIRR cells even though they did lengthen the 3′ UTR of *MDM2* (Figure S4G).

Considering the essential role of the IRR of *MDM2* in regulating the p53 signaling pathway, we examined its interspecies conservation. Considering the length of 3′ UTR of human *MDM2* is ∼11 kb, we performed comparative genomic analysis using 15 kb sequences from the stop codon of *MDM2* mRNA of various primate species as well as of some murine species (Figure 5J). We found that the sequence similarities showed great concordance with the phylogenetic groups. For example, the *MDM2* 3′ UTR of primates closely related to humans (Catarrhini) showed sequence similarities of over 90%, while *the MDM2* 3′ UTR of New World monkeys (Platyrrhini) showed sequence similarities ranging from 80% to 90% to the 3′ UTR of human *MDM2*. We next examined the status of IRAlus insertion at the 3′ UTR of *MDM2* using RepeatMasker. Interestingly, IRAlus elements of the *MDM2* 3′ UTR were highly conserved in the distal position of the 3′ UTR, especially in apes (Figure 5J). More importantly, all examined species showed heavy Alu clusters in the putative aUTRs of *MDM2.* Even in murine species, despite relatively low sequence similarity, we could find inverse SINE/B1 and B2 elements in the distal UTR region. This indicates that Alus, especially IRAlus or inverted SINEs, may serve an important regulatory function, at least for *MDM2*.

### Regulation of p53 pathway by MDM2 UTR shortening in lung adenocarcinomas

To extend our findings and test the clinical relevance, we reanalyzed publicly available RNA-seq data generated from fresh surgical specimens of human lung adenocarcinoma tumors and their adjacent normal tissues.^32^ First, we confirmed the upregulation of *CSTF2* in lung adenocarcinomas (Figures 6A and S5A). While 18 out of 69 pairs showed a more than 2-fold increase in *CSTF2* expression, only one tumor showed a 2-fold downregulation in *CSTF2* expression (Figure S5A). In addition, we found lung tumors with high *CSTF2* expression exhibited the 3′ UTR shortening event of *MDM2* (Figure 6B). Moreover, GSEA analysis clearly showed the repression of the p53 pathway in lung tumors with high *CSTF2* expression, which is consistent with our *in vitro* data of p53 activation in *CSTF2* and *CSTF2T* deficient A549 cells (Figures 6C and S5B). Interestingly, most of the significantly enriched Hallmark gene sets identified by GSEA overlapped with those affected by the knockdown of *CSTF2* and *CSTF2T*, such as TNF-α signaling via NF-κB, inflammatory response, and apoptosis (Figure 6C). Analysis of p53 target genes further confirmed the repression of the p53 pathway in lung tumors with high *CSTF2* expression (Figure 6D). Correlation analyses with a total of 69 tumor/normal lung adenocarcinoma pairs also supported our *in vitro* data (Figures 6E and S5C). Collectively, our analysis suggests that tumors might utilize global UTR shortening to exploit the gene-silencing effect of 3′ UTR IRAlus of *MDM2* to suppress p53 activation.

**Figure 6.**
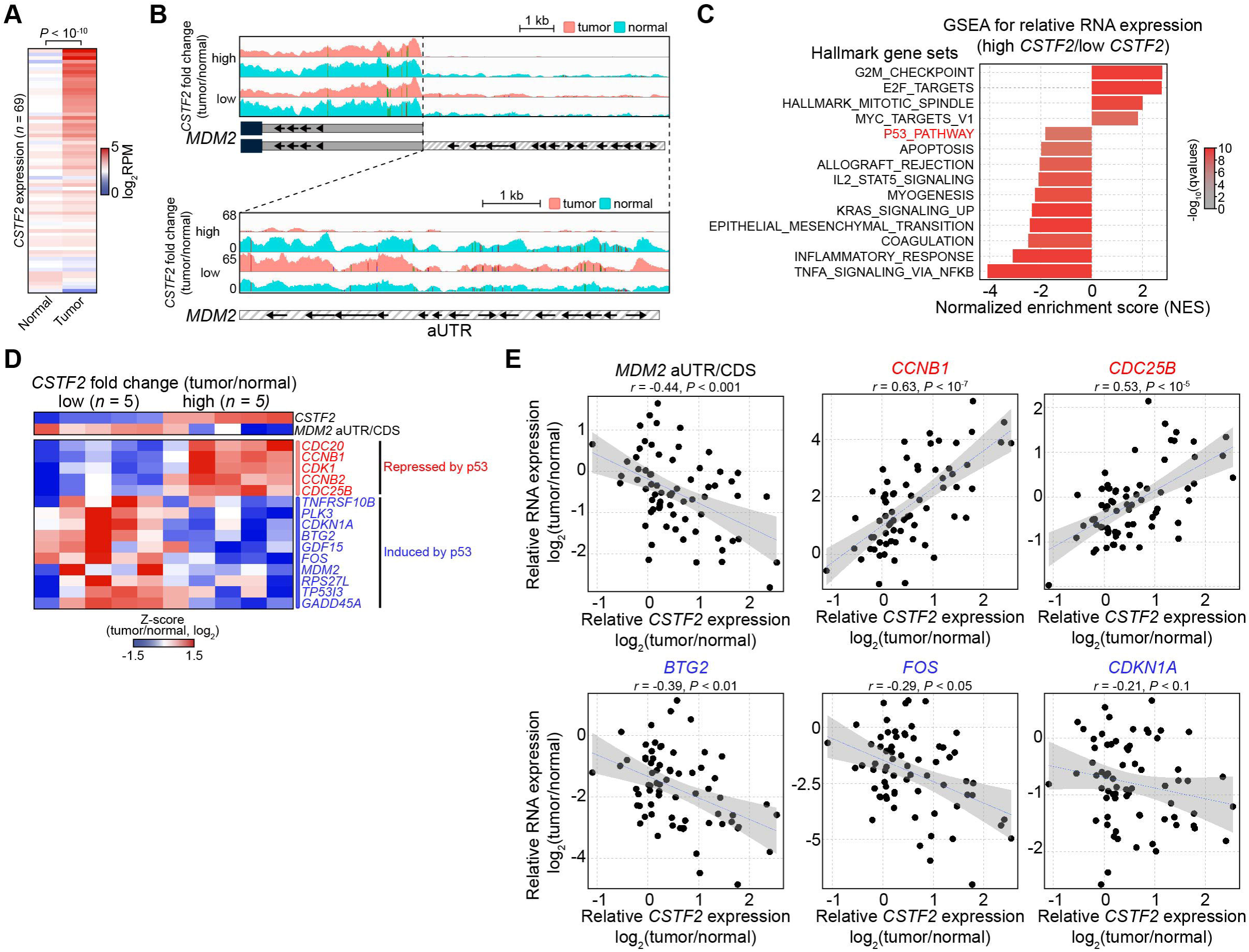
Regulation of the p53 pathway in lung adenocarcinomas. (A) *CSTF2* mRNA expression in human lung adenocarcinomas and their adjacent matched-normal tissues (*n* = 69). (B) IGV snapshot of RNA-seq reads at MDM2 3′ UTR in human lung adenocarcinoma tissues with indicated relative expression of *CSTF2*. (C) Significantly enriched Hallmark gene sets identified by GSEA for the relative gene expression in human lung adenocarcinomas with high *CSTF2* expression (*n* = 3). (D) Heatmap of APA level of *MDM2* (*MDM2* aUTR/CDS) and relative expression of genes previously identified as directly induced or repressed by the p53 pathway in human lung adenocarcinoma tissues with the indicated relative expression of *CSTF2*. (E) Correlation between relative *CSTF2* expression and APA level of *MDM2* or relative expression of genes from B (repressed by p53: *CCNB1*, *CDC25B*; induced by p53: *BTG2*, *FOS*, *CDKN1A*) in human lung adenocarcinoma tissues (*n* = 69). Correlation *r* and its *P* was calculated using the Spearman correlation.

### Neuronal cell-specific gene silencing by IRAlus

Encouraged by these results, we further investigated cell-specific gene regulation by IRAlus during global APA. One important example of tissue-specific APA change is transcriptome-wide 3′ UTR lengthening in neuronal cells. Both microarray analysis and deep sequencing showed the global 3′ UTR lengthening in the mammalian brain.^33,34^ Moreover, the central nervous system (CNS)-specific 3′ UTR lengthening occurred in flies,^35^ supporting the conserved phenomenon of UTR lengthening in the nervous system. In this context, we investigated whether long 3′ UTRs of genes expressed neuronal cells were more susceptible to IRAlus-mediated gene silencing.

We first confirmed the expression of two isoforms of *NEAT1* RNA (*NEAT1_1* and *NEAT1_2*) in NPCs by reanalyzing nuclear and cytosolic total RNA-seq data (Figure S6A).^36^ Next, we performed J2 fCLIP-seq on NPCs derived from human embryonic stem cells (hESCs) and examined the J2 enrichment pattern. The overall pattern was similar to that of HeLa cells, but we found a dramatic increase in the nuclear retention of conditional and extended 3′ UTR IRAlus genes (Figures 7A and 7B), consistent with the global 3′ UTR lengthening in these cells. Of note, to consider the 3′ UTR isoforms longer than the known 3′ UTR annotation, we extended the 3′ UTRs by up to 2,000 mer when calculating the J2 enrichment scores. Moreover, to compare the J2 enrichment score and nuclear retention levels of mRNAs between HeLa and NPCs, those values were normalized to the median value of noIRAlus genes. NPCs also showed J2 enrichment score-dependent nuclear retention of mRNAs with stronger significance than that of HeLa cells (Figure 7C). Closer inspection revealed strong J2 enrichment of the aUTR of conditional IRAlus genes as well as conditional (*MXRA7* and *LLPH;* Figure 7E) and extended 3′ UTR IRAlus (*C11orf1*; Figure 7F). Altered J2 enrichment due to 3′ UTR isoform switching (*RPL15*; Figure S6B) was also apparent in NPCs.

**Figure 7.**
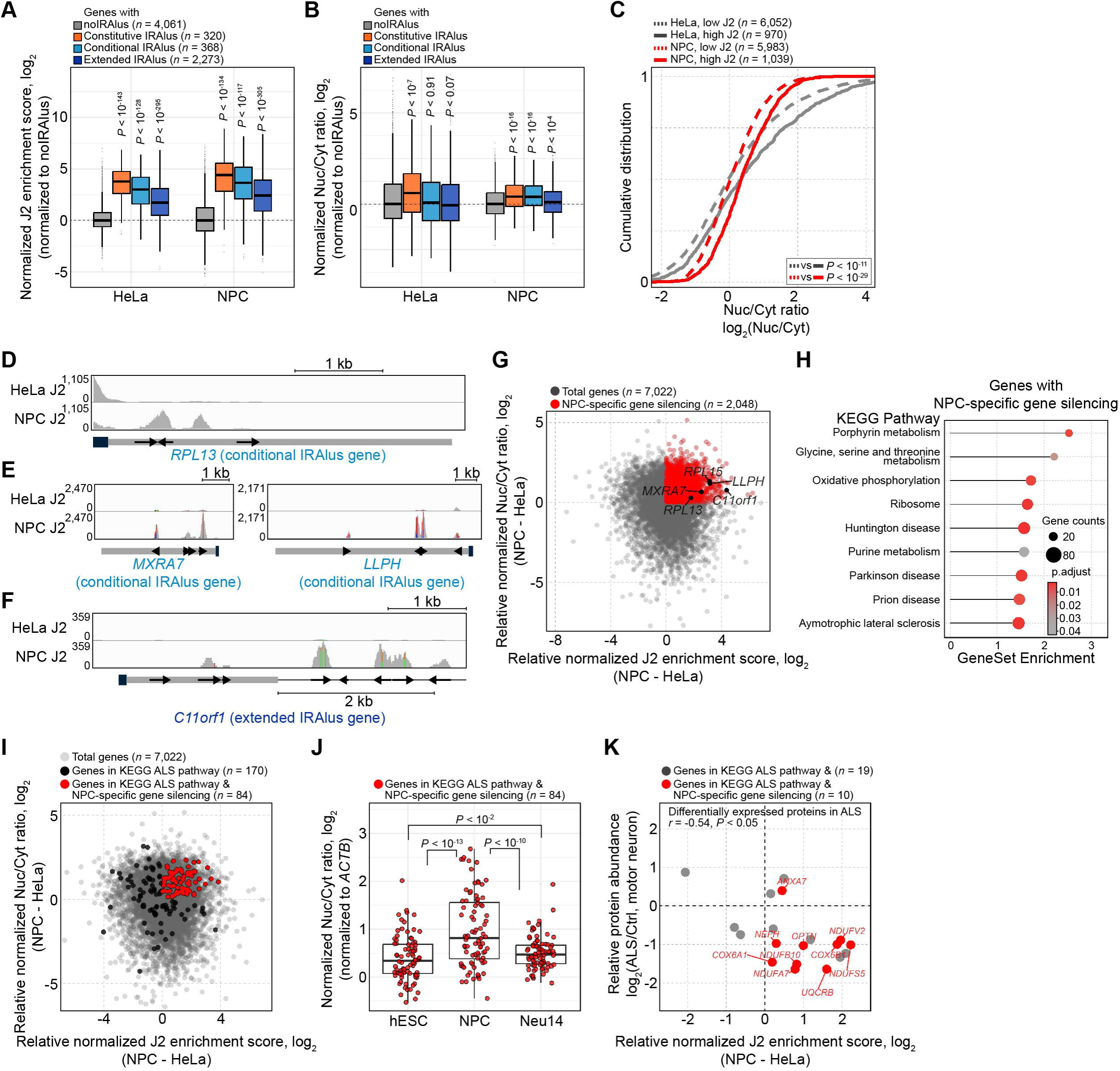
NPC-specific gene regulation by IRAlus. (A and B) Normalized J2 score (A) and Nuc/Cyt ratio (B) of mRNAs with the indicated 3′ UTR IRAlus in HeLa and NPCs. (C) Cumulative distribution of the Nuc/Cyt ratio of mRNAs with the indicated J2 scores. (D-F) IGV snapshot of J2 fCLIP-seq reads in HeLa and NPC cells for selected conditional (D, *RPL13;* E, *MXRA7* and *LLPH*) or extended (F, *C11orf1*) IRAlus gene. (G) Scatter plot of relative normalized J2 score and Nuc/Cyt ratio of mRNAs in NPCs when compared to those in HeLa. Genes with NPC-specific gene silencing effect are indicated in red (*n* = 2,048). NPC-specific gene silencing effect was defined as a log_2_ fold change of normalized J2 score > 0 & log_2_ fold change of normalized Nuc/Cyt ratio > 0. (H) Over-representation analysis of genes showing NPC-specific gene silencing effect in the KEGG pathway using clusterProfiler. (I) Scatter plot of relative normalized J2 score and Nuc/Cyt ratio of mRNAs in NPCs when compared to those in HeLa. Genes involved in the KEGG ALS pathway are indicated in black (*n =* 170), and those associated with the ALS pathway with increased J2 and Nuc/Cyt ratio are marked in red (*n* = 84). (J) Normalized Nuc/Cyt ratio of 84 genes marked as red in (I) in hESCs, NPCs, and Neu14. (K) Scatter plot of relative protein abundance of differentially expressed proteins (*n* = 19) of the spinal cord of patients with ALS by referring to the previous study. Genes involved in the KEGG ALS pathway with NPC-specific gene silencing are indicated in red (*n* = 10). All *P* were calculated using the two-sided Wilcoxon signed rank test.

We then identified genes showing both high J2 enrichment scores and nuclear retention in NPCs when compared to those of HeLa cells (*n* = 2,048; Figure 7G) and subjected them to the Kyoto Encyclopedia of Genes and Genomes (KEGG) pathway enrichment analysis. Interestingly, these NPS-specific IRAlus genes were strongly associated with neurodegenerative disorders, such as amyotrophic lateral sclerosis (ALS), Parkinson’s disease (PD), and Huntington’s disease (HD; Figure 7H). We focused our analysis on ALS as RNA and protein mislocalization were closely associated with the pathogenesis of ALS.^37,38^ We found that out of 170 genes involved in the KEGG ALS pathway, 84 (49.4%) showed increased J2 enrichment scores as well as increased Nuc/Cyt ratios in NPCs compared to those of HeLa cells (Figure 7I). Interestingly, these 84 genes showed decreased Nuc/Cyt ratios in 14-day differentiated neurons (Neu14)^36^ as these cells did not exhibit nuclear paraspeckles due to a lack of nuclear *NEAT1* lncRNA expression (Figure 7J).^39,40^ Of note, hESCs were used as a negative control as these cells also lack *NEAT1* RNA and do not have paraspeckles.^8^ Indeed, the upregulation of *NEAT1* expression and subsequent paraspeckle formation in the motor neurons of ALS patients^39,40^ further supported our hypothesis that misregulation of 3′ UTR IRAlus genes due to an increased level of *NEAT1* resulted in the silencing of genes involved in ALS. Consistent with this idea, when we analyzed the protein expression by referring to the previous proteomics study,^41^ we found that out of 19 ALS-associated genes, ten of them (52.6%) showed increased J2 enrichment scores and Nuc/Cyt ratios in NPCs compared to those of HeLa cells and nine of these ten showed decreased protein expression in motor neurons of ALS patients (Figure 7K).

We further investigated the NPC-specific IRAlus genes and found that the majority of them were nuclear-encoded mitochondrial protein-coding genes. Gene ontology (GO) over-representation analysis of genes involved in ALS showed a clear relevance to mitochondrial functions, including ATP synthesis and mitochondrial respiratory chain complex (RCC; Figures S6C and S6D). Closer inspection of J2 fCLIP-seq in NPCs revealed that these genes involved in mitochondrial functions (NDUFS5: NADH; COX6B1, UQCRB: cytochrome *c*) mostly contain IRAlus elements in their extended 3′ UTR (Figures S6E-G). This is consistent with the recently reported association between mitochondrial dysfunction and ALS.^42^ Moreover, GO analysis also revealed the enrichment of mRNA transport-related genes, further suggesting the importance of proper regulation of 3′ UTR IRAlus genes during ALS pathogenesis (Figure S6C). Altogether, our data indicate that 3′ UTR lengthening in neuronal cells is linked to IRAlus-mediated brain-specific gene regulation. Additionally, with aberrant *NEAT1* lncRNA expression and paraspeckle assembly, this brain-specific gene regulation may play an important role in the progression of neurodegenerative disorders, especially ALS.

## DISCUSSION

In this study, we directly captured and mapped the dsRNA structure generated by the IRAlus element using the J2 antibody and identified genes that were under the regulation of the 3′ UTR IRAlus. Through such approach, we showed that (1) many genes contain Alu elements in aUTRs or even downstream of the annotated 3′ end, thus having a potential to be regulated by IRAlus-mediated gene silencing during APA; (2) Global 3′ UTR lengthening resulted in over a hundred genes to be regulated by 3′ UTR IRAlus, including *MDM2*; (3) IRR in the aUTR of *MDM2* mRNA could regulate the RNA localization and protein expression and subsequently affect the activation status of p53; (4) UTR shortening during tumorigenesis may suppress p53 signaling by relieving IRAlus-mediated gene silencing of MDM2; (5) NPC-specific IRAlus genes were associated with neurodegenerative diseases, including ALS; (6) Many of the genes associated with ALS pathogenesis contained IRAlus in their 3′ UTR and were downregulated in the spinal cord of ALS patients. Collectively, our study establishes that hundreds of genes have the potential to be regulated by 3′ UTR IRAlus during APA and that the proper regulation of 3′ UTR IRAlus is closely associated with human pathophysiology.

By directly capturing dsRNA structures, we could accurately identify genes under the regulation of 3′ UTR IRAlus without the need for information on UTR variants. Moreover, our list of genes shows significant nuclear sequestration and translational suppression. Notably, most of the genes contained IRAlus elements either in the aUTR (long UTR) variant or even downstream of the annotated 3′ end. The genes with IRAlus in the major UTR variant only occupied about 43% of all genes with high J2 enrichment scores. In addition, most of the genes with high J2 enrichment scores but without IRAlus element contained at least one Alu element in their 3′ UTRs, suggesting their potential regulation by another Alu element *in trans* or by another Alu element downstream of the annotated 3′ end. Alternatively, these could reflect a single Alu element that retains the mRNA in the nucleus.^43^ These results indicate that IRAlus may induce cell-specific/tissue-specific gene regulation that accompanies changes in the 3′ UTR length. Profiling of genes with 3′ UTR IRAlus element using J2 fCLIP-seq will enhance our understanding of the regulatory potential of the IRAlus element in shaping tissue-specific gene expression landscape.

Our study suggests that global 3′ UTR shortening may contribute to tumorigenesis through the translational activation of *MDM2* via IRAlus exclusion and subsequent repression of the p53 pathway. Indeed, we found a negative correlation between *CSTF2* expression and distal poly(A) site usage of *MDM2* and p53 pathway activation in both cells and lung adenocarcinomas. Moreover, our *in vitro* experiments using lung adenocarcinoma cells showed the importance of IRR in the aUTR in modulating *MDM2* mRNA nuclear sequestration and downregulation of MDM2 protein expression during UTR lengthening. The striking interspecies conservation of the Alu cluster in the *MDM2* 3′ UTR further suggested the likely functional significance of IRR. Considering that inducing UTR lengthening of *MDM2* mRNA using ASOs resulted in increased p53 activation and cell death, future investigation and optimization of *MDM2* targeting ASOs may present novel therapeutic strategies targeting wild-type p53 cancers with increased CSTF2 expression by reactivating the p53 pathway.

One of the important findings of this study is that global UTR lengthening in NPCs shapes neuronal cell-specific post-transcriptional regulation by 3′ UTR IRAlus. We found that most of the genes showing neuronal cell-specific gene silencing were involved in the progression of neurodegenerative disorders. Interestingly, previous studies showed that normal postmitotic neurons were devoid of nuclear paraspeckles due to mislocalization and low expression of *NEAT1* lncRNA,^39,40,44^ indicating that neurons avoid IRAlus-mediated gene silencing by controlling *NEAT1*. Consistent with this idea, recent studies have demonstrated a strong association between neurodegenerative disorders and aberrant hyper-assembly of paraspeckles through altered *NEAT1*_2 expression.^39,40^ This increase in paraspeckle formation was believed to be one of the defense mechanisms against stress conditions.^40^ Our study suggests otherwise that the elevated *NEAT1_2* lncRNA may induce 3′ UTR IRAlus-mediated gene silencing effects globally, resulting in the disruption of mRNA transport and mitochondrial metabolism. The potential of *NEAT1_2* RNA being the master regulator of global 3′ UTR IRAlus regulation during ALS pathogenesis needs further investigation in the future.

In summary, our transcriptomic analysis, followed by functional validation, highlights the crosstalk among APA, IRAlus, and paraspeckles in shaping the cell-specific gene regulatory landscape. Moreover, our findings provide an additional layer of post-transcriptional gene regulation and underline the need for further investigation on biological processes and pathogenesis of diseases that accompany global APA change or aberrant paraspeckle formation.

### Limitations of the study

When identifying the IRAlus element, additional possible parameters, such as the length or subtype of Alu and the presence of other adjacent *cis*-elements, could be considered. With well-controlled massive screening using constructed libraries, future comprehensive studies could provide a better understanding and further application of IRAlus-mediated gene silencing. Next, the potential contribution of the 3′ UTR IRAlus and aberrant formation of paraspeckle in neurodegenerative disorders relied on J2 fCLIP-seq performed in NPCs. Owing to a large amount of total RNA requirement, we could only perform J2 fCLIP-seq in NPCs rather than in neurons. We anticipate that technological advances will overcome this hurdle and provide direct evidence of enhanced gene silencing in neurons. Lastly, we only analyzed the role of global APA changes by CSTF2 during tumorigenesis. With the diverse polyadenylation factors involved in global APA changes during tumorigenesis, comprehensive studies will be required to uncover the biological role of IRAlus in pathological conditions that accompany global APA change.

## ACKNOWLEDGEMENTS

We thank all the members of the Yoosik Kim laboratory for their helpful discussions and comments on the paper. We thank Professor Jinkuk Kim from KAIST for his help with designing ASOs. This study was supported by the Basic Research Laboratory Program through the National Research Foundation of Korea (grant number NRF-2021R1A4A3032789), funded by the Korean government’s Ministry of Science and ICT. We also acknowledge KARA (KAIST Analysis Center for Research Advancement) for providing the experimental equipment.

## AUTHOR CONTRIBUTIONS

J. Ku, J. Han, Y-s. Lee, and Y. Kim designed the study and analysis. Experiments were performed by J. Ku. S. Kim, K. Lee, D. Ku, and N. Kim. J. Ku and H. Lee performed bioinformatic analysis. H. Do prepared NPCs. The study was supervised by Y. Kim. J. Ku and Y. Kim wrote the manuscript with contributions from S. Kim, H. Do, J. Han, and Y-s. Lee.

## RESOURCE AVAILABILITY

### Lead contact

Further information and requests for resources and reagents should be directed to and will be fulfilled by the lead contact Yoosik Kim.

### Materials availability

All reagents and resources used for this study are available upon request to the lead contact.

### Data and code availability

- J2 fCLIP-seq and mRNA-seq following subcellular fractionation data have been deposited in GEO and are publicly available as of the date of publication. Accession numbers are listed in the key resources table. Original western blot images, gel images, and microscopy data have been deposited at Mendeley and are publicly available as of the data of publication. The DOI is listed in the key resources table.
- No new software was developed during this study.
- Any additional information required to reanalyze the data reported in this paper is available from the lead contact upon request.

### EXPERIMENTAL MODELS AND SUBJECT DETAILS

#### Cell lines and culture

HeLa and HEK-293T cells were grown in Dulbecco’s modified Eagle medium (DMEM; Welgene) supplemented with 9.1% fetal bovine serum (FBS; Gibco). A549 cells were grown in RPMI medium (Welgene) supplemented with 9.1% FBS. Cells were grown in a 37 °C and 5% CO_2_ humidified incubator and passaged at a 1:5 dilution every 2-3 days. H9 (WA09) human embryonic stem cells (hESCs) were cultured on mitomycin C (AG Scientific) treated mouse embryonic fibroblasts (MEF) in hESC medium. The hESC medium consisted of DMEM/F12 (Gibco) with 1% MEM non-essential amino acids (NEAA) (Gibco), 1.2 mM sodium bicarbonate (Sigma Aldrich), 1 mM L-glutamine (SAFC), 0.1 mM 2-mercaptoethanol (Sigma Aldrich), 20% of KnockOut serum replacement (Gibco), and 10 ng/ml bFGF (R&D systems). The cells were passaged using 10 μg/ml collagenase IV (Gibco). To generate embryoid bodies (EBs), hESC colonies were dissociated and plated on non-adherent dishes. After one day of floating culture, the EBs were treated with 0.2 μM LDN (Selleckchem) and 10 μM SB-431542 (Cayman Chemical) in DMEM/F12□+□GlutaMAX (Gibco) plus N-2 and B-27 supplements (Gibco). The treatment was continued for 7 days, and the EBs were plated on growth factor reduced matrigel (Corning)-coated dishes in DMEM/F12 plus N-2 and B-27 supplements (N2B27 medium) 0.2 μM LDN, 10 μM SB-431542, and 1 μg/ml laminin (Gibco). Neural rosettes were manually collected and dissociated with Accutase (Innovative Cell Technologies) and plated onto poly-L-ornithine (Sigma Aldrich)/laminin-coated dishes with NPC medium (N2B27 medium plus 20 ng/ml bFGF). Protocols describing the use of human ESCs were approved in accordance with the ethical requirements and regulations of the Institutional Review Board of KAIST (IRB #KH2021-069).

### METHOD DETAILS

#### Chemical treatment and cell viability test

siRNAs were transfected to cells using Lipofectamine 3000 (Invitrogen) following the manufacturer’s protocol. To knockdown the expression of *NEAT1* lncRNA, Lipofectamine RNAiMAX (Invitrogen) was used. For the ASO transfection, Lipofectamine RNAiMAX (Invitrogen) was used. siRNAs and ASOs were purchased from Bioneer Korea and IDT, respectively. The sequence of non-target ASO was referred from a previous study.^31^ The sequences of the siRNAs and ASOs used in this study are summarized in Table S1. To block proteasome, 10 μM of MG132 (Sigma Aldrich) was added to the medium. For the cell viability test, the Cell Counting Kit-8 (CCK-8; Dojindo) was used following the manufacturer’s protocol.

#### Subcellular fractionation

To fractionate cells into nuclear and cytosolic fractions, the NE-PER Nuclear and Cytoplasmic Extract Reagent kit (Thermo Scientific) was used. Cells were subsequently lysed with CER I and CER II buffers at 4°C for 10 min and 1 min, respectively. The cells were centrifuged at 20,000 × g at 4°C for 5 min to obtain the cytosolic fraction (supernatant). The pellet fraction was washed twice with phosphate buffer saline (PBS) and further lysed with the NER buffer at 4°C for 40 min. The nuclear fractions were then obtained by centrifuging the lysates at 20,000 × *g* at 4°C for 10 min.

#### RNA extraction and reverse transcription-quantitative PCR (RT-qPCR)

Total RNA was extracted from cells or fractionated lysates using TRIzol (Invitrogen) or TRIzol LS (Invitrogen), respectively. Extracted nucleic acids were treated with DNase I (Takara) and reverse-transcribed using RevertAid reverse transcriptase (Thermo Scientific) or SuperScript IV reverse transcriptase (Invitrogen) using random hexamer primers (Invitrogen). The synthesized cDNA was amplified using the SensiFAST SYBR Lo-Rox Kit (Bioline) and analyzed using an AriaMX Real-Time PCR System (Agilent) or QuantStudio 1 (Applied Biosystems). For multiplex RT-PCR, the synthesized cDNA was PCR amplified using the KOD-Plus-Neo (TOYOBO), and the PCR amplicon was analyzed on a 3% TBE agarose gel. The sequences of primers for RT-qPCR and multiplex RT-PCR used in this study are summarized in Table S2.

#### RNA-fluorescent in situ hybridization (RNA-FISH)

RNA-FISH probes were prepared using the MEGAscript T7 Transcription Kit (Invitrogen) and DIG RNA Labeling Mix (Roche). The DNA template for *in vitro* transcription was amplified by KOD-Plus-Neo (TOYOBO) using cDNA and reverse primer with the T7 promoter sequences at the 5′ termini. *In vitro* transcription was performed using a DIG labeling mix (Roche) at 37°C overnight. Template DNA was removed by Turbo DNase I digestion at 37°C for 15 min. The RNA probe was purified by acid phenol:chloroform extraction and ethanol precipitation. The purified probe was dissolved in 20 μl of hydrolysis buffer (40 mM NaHCO_3_ and 60 mM Na_2_CO_3_, pH 10) and fragmented at 60°C for 40 min. Hydrolysis was stopped by adding an equal volume of stop buffer (0.2 M NaOAc, pH 6), and the probe was immediately quenched on ice. The RNA probe was then diluted in 200 μl of hybridization buffer (50% formamide, 10% dextran sulfate, 0.1% SDS, 300 ng/mL salmon sperm DNA (Sigma Aldrich), 2X SSC, and 2 mM vanadyl ribonucleoside complex).

Cells cultured on a confocal dish were fixed in 4% formaldehyde (Thermo Scientific) for 10 min at room temperature. Fixed cells were permeabilized in (70% ethanol, 2X SSC) overnight at 4°C. The cells were rehydrated by successive washing with (50% ethanol, 2X SSC) and (25% ethanol, 2X SSC) for 3 min each at room temperature. Cells were incubated in (50% formamide, 2X SSC) at 40°C for 2 h and hybridized at 40°C overnight with an RNA probe. Before hybridization, the RNA probe was denatured by heating at 85 °C for 5 min and quenched on ice for at least 1 min. Hybridized cells were stringently washed twice with (50% formamide, 2X SSC) and (50% formamide, 0.1X SSC) at 40°C for 30 min each. The cells were blocked in a blocking buffer (0.2% BSA, 8% formamide, 2X SSC) at room temperature for 1 h. Cells were incubated with anti-DIG antibody (Roche), 1:100 diluted in blocking buffer at room temperature for 2 h. Alexa Fluor-conjugated secondary antibody (Invitrogen) was used to label the primary antibody. Cells were also incubated with DAPI (Invitrogen) solution to counterstain the nuclei. The stained cells were imaged with a Zeiss LSM780 or LSM880 confocal microscope using a 63X objective. The sequences of the PCR primers used for *in vitro* transcription of the template DNA used in this study are summarized in Table S3.

To calculate the colocalization rate with DAPI, the blue channel was used to detect the nucleus. For each image, connected pixels of the blue channel were identified and considered as the nucleus. The colocalization rate was calculated by dividing the intensity of the red channel (RNA detection channel) colocalized in the nucleus by that of the total intensity of the red channel.

#### Lentivirus production and transduction

Lentivirus was produced according to published protocols,^45,46^ with some modifications. HEK- 293T cells were prepared at 70% confluence in a 100-mm dish with 5 ml of medium. Lentivirus packaging plasmids (7.5 μg of transfer plasmid, 5.6 μg of psPAX2, and 3.75 μg of pMD2.G) were transfected using Lipofectamine 3000 (Invitrogen). After 6 h, the medium was replaced with fresh DMEM with 9.1% FBS (Gibco) supplemented with 1% BSA to improve virus stability. After 60 h, the viral supernatants were harvested and centrifuged at 3,000 rpm at 4°C for 10 min to pellet the cell debris. The supernatant was then filtered through a 0.45 μm SFCA syringe filter (Corning) and stored at -80°C.

To transduce the lentivirus, the cell medium was replaced with lentivirus medium 1:4 diluted in fresh medium supplemented with 10 μg/ml of polybrene (Sigma Aldrich). After 2 days, the cell medium was replaced with fresh medium, and 10 μg/ml of blasticidin (Gibco) or puromycin (InvivoGen) was added every 2 days until the transduction control cells died. After selection, cells were grown without blasticidin or puromycin for additional 2 days before further analysis.

#### Gene deletion using CRISPR-Cas9

To generate A549 cells stably expressing Cas9, a lentivirus was generated using lentiCas9-Blast (Addgene) and transduced into A549 cells. Single-cell clones of A549-Cas9 were selected after blasticidin (Gibco) treatment and expanded in a 96-well plate. To delete IRAlus in the MDM2 aUTR, a total of four sgRNAs, two targeting the upstream and the other two targeting the downstream of the IRR of MDM2 aUTR, were designed and inserted into the lentiGuide-Puro vector individually using BsmBI restriction sites. A mixture of lentiviruses of the four sgRNAs was transduced into A549-Cas9 cells. Single-cell clones of A549-ΔIRR were selected after puromycin treatment and expanded. Deletion of gDNA was confirmed by PCR with gDNA extracted with the Quick-DNA Miniprep Plus Kit (Zymo Research) using KOD-Plus-Neo DNA polymerase (TOYOBO). The PCR product was further analyzed by Sanger sequencing. The sequences of sgRNAs are summarized in Table S4. The sequences of primers for gDNA amplification are summarized in Table S5.

#### Western blotting

Total cell lysates were prepared by sonication in lysis buffer (50 mM Tris-HCl pH 8.0, 100 mM KCl, 0.5% NP-40, 10% glycerol, and 1 mM DTT) supplemented with protease inhibitor cocktail III (Calbiochem) and phosphate inhibitor cocktail I (AG Scientific). A total of 40–50 μg of protein was separated on an SDS-PAGE gel and transferred to a polyvinylidene difluoride (PVDF) membrane using an Amersham semi-dry transfer system. Following primary antibodies were used: TUBB (86298), Rab5 (3547), p-p53 (9284), MDM2 (86934), and CDKN1A (2947) were purchased from Cell Signaling Technology; GAPDH (sc-32233) and Lamin A/C (sc-376248) were purchased from Santa Cruz Biotechnology. All antibodies were used at 1:1000 dilution.

#### J2 formaldehyde-mediated crosslinking immunoprecipitation

To prepare J2 antibody-conjugated resin, Pierce Protein A UltraLink Resin (Thermo Scientific) was incubated with J2 antibody (Jena Bioscience) in the fCLIP lysis buffer (20 mM Tris-HCl pH 7.5, 15 mM NaCl, 10 mM EDTA, 0.5% NP-40, 0.1% Triton X-100, 0.1% SDS, 0.1% sodium deoxycholate) on a rotator at 4°C for 3 h. The harvested cells were fixed with 0.1% formaldehyde (Thermo Scientific) for 10 min at room temperature. Cells were immediately quenched by adding 1/3 volume of 1 M glycine and incubated at room temperature for an additional 10 min. Crosslinked cells were lysed in the fCLIP lysis buffer on ice for 10 min, and subsequently sonicated multiple times for complete lysis. The lysate was then supplemented with 50 U/ml (12 nM) of RNase T1 (Worthington) and immunoprecipitated using J2 antibody-conjugated resin in a rotator at 4°C for 3 h. The J2 antibody-conjugated resin was washed with fCLIP wash buffer (20 mM Tris-HCl pH 7.5, 150 mM NaCl, 10 mM EDTA, 0.1% NP-40, 0.1% Triton X-100, 0.1% SDS, 0.1% sodium deoxycholate). Immunoprecipitated RNAs were eluted using urea elution buffer (200 mM Tris-HCl pH 7.5, 100 mM NaCl, 20 mM EDTA, 2% SDS, and 7 M Urea) at 25°C with 950 rpm for 3 h in a ThermoMixer (Eppendorf). The eluted RNA was then de-crosslinked and separated from the J2 antibody and other proteins by incubating at 65°C at 950 rpm overnight with 2 mg/ml Proteinase K (Roche) in a ThermoMixer. RNA was purified using acid phenol:chloroform extraction and DNA contaminants were digested using DNase I (Takara).

#### Identification of 3′ UTR IRAlus genes

A list of 3′ UTR IRAlus genes was generated by analyzing the most recent version of the genomic annotation for Alu elements (from RepeatMasker) and transcriptome information (from GENCODE). There was no favorable Alu population to be located in 3′ UTRs as 3′ UTR Alus were indistinguishable from Alus in other locations (Figure S7A). The distance between two inverted Alu elements (IRAlus distance) was optimized by comparing the Nuc/Cyt ratio of mRNAs of IRAlus genes with that of noIRAlus genes. Genes with a long IRAlus distance (longer than 2,500 nt) still resulted in the nuclear retention of the host mRNA (Figure S7B). Hence, an IRAlus distance of 6,000 nt (93^rd^ percentile of all IRAlus distances) was used as an upper limit for IRAlus gene identification. With this limit, more than 6,500 genes express 3′ UTRs that were long enough to potentially contain IRAlus elements with a median IRAlus distance of 1,557 nt (Figure S7C). Using this optimized IRAlus annotation, 699 IRAlus-containing genes (8.5%) out of 8,243 abundantly expressed genes in HeLa cells were identified (Figure S7D).

#### Characterization of 3′ UTR IRAlus genes

Alu and IRAlus elements located in the 3′ UTR were identified using transcriptome information from GENCODE v27^47^ and Pre-Masked RepeatMasker for hg38. Only 3′ UTR IRAlus with a distance between two inverted Alu elements of less than 6,000 nt were considered for further analyses. For IRAlus gene identification, the number of IRAlus located in all annotated 3′ UTRs of a given gene was determined. If the gene has at least one IRAlus in its 3′ UTR, it was considered an IRAlus gene. For the IRAlus gene identification with APA, the major 3′ UTR was defined as the 3′ UTR isoform produced by cleavage at the strongest poly(A) site based on the PolyASite 2.0 database^19^ with the following criteria: (average expression > 0.05, percentage of samples > 0.05) or poly(A) site with the highest average expression. If a gene did not have poly(A) site annotation in PolyASite 2.0 database, the 3′ end of the longest 3′ UTR of the gene was considered as the major 3′ UTR. If a gene had at least one IRAlus element in its major 3′ UTR, the gene was considered a “constitutive IRAlus gene”. All other IRAlus genes were considered to be “conditional IRAlus genes”. In addition, to consider the genes containing IRAlus elements in its extended 3′ UTR, genes containing at least one IRAlus element in the region up to 2,000 mer before reaching a neighboring gene from the annotated 3′ ends were determined and considered as “extended IRAlus genes”.

#### RNA-seq library preparation and alignment to the genome

2×101 paired-end total RNA-seq and mRNA-seq libraries were constructed using the TruSeq Stranded Total RNA with Ribo-zero H/M/R Gold kit (Illumina) and TruSeq Stranded mRNA Sample Prep Kit (Illumina), respectively. The RNA-seq libraries were sequenced using NovaSeq 6000 platform (Illumina). Raw paired-end sequencing reads were mapped to the reference human genome (build hg38) using HISAT2 v.2.1.0^48^ using default parameters, except with the option “--rna-strandness RF”. Uniquely and concordantly mapped reads were selected using samtools^49^ and then were used for further analyses.

#### Annotation of RNA-seq alignments and data processing

To assign and summarize genomic features, the featureCounts function of the Rsubread v2.8.2^50^ R package was used with transcriptome information from GENCODE v27 or RepeatMasker pre-masked for hg38. For the GENCODE, only protein-coding transcripts with a transcript_support_level of 1 or 2 were used. To calculate the Nuc/Cyt ratio, translational efficiency, and J2 enrichment score, raw read counts normalized to reads per million (RPM) were used. The DESeq2 v1.34.0^51^ R package was used to analyze differentially expressed genes (DEGs) with summarized raw read counts for each gene. For GSEA and over-representation analyses, the clusterProfiler v4.2.2^52^ R package and DOSE v.3.20.1^53^ R package were used. Hallmark gene sets were obtained from the Molecular Signatures Database (MSigDB) v7.4.^23^ For the secondary structure prediction of RNAs, RNAfold web server^54^ was used.

#### Analysis of miRNA binding sites

Abundance estimates of miRNAs in HeLa cells were obtained from DIANA-miTED.^18^ 22 miRNAs with an abundance of > 1% of the overall miRNA were chosen. Their probability of conserved targeting (Aggregate P_CT_ ^16^) and targeting efficacy (CWCS^17^) were obtained from the TargetScan 8 database.^55^ Information on half-lives of mRNAs in HeLa cells was obtained from a previous study.^56^

#### Interspecies conservation of MDM2 3′ UTR

Sequences of the putative 3′ UTR of the MDM2 gene (15,000 nt downstream of the stop codon) of several primates and murine were aligned and their sequence identities were analyzed using Clustal Omega.^57^ Embedded repeat elements (Alu or Alu-like SINE/B1) in the putative 3′ UTR of the MDM2 gene of each species were identified using RepeatMasker v4.1.4 and Dfam v3.6.

#### Lung cancer and matched tissue RNA-seq analysis

Raw mRNA-seq data of 69 pairs of lung adenocarcinomas and their adjacent normal tissues were obtained from a previous study.^32^ RNA fold change (tumor/normal) of genes was measured and used for further analyses. For the MDM2 APA level calculation, the sum of read counts aligned to the following genomic region was used: MDM2 CDS: chr12:68839274- 68839849; MDM2 aUTR: chr12:68844193-68850686.

#### Analysis of protein abundance in spinal cord tissues of patients with ALS

The abundance of differentially expressed proteins in the spinal cord tissues of patients with ALS was obtained from a previous study.^41^ Among a total of 293 proteins, proteins involved in ALS as well as proteins showing NPC-specific gene silencing were identified and compared.

#### Quantification and statistical analysis

Statistical analyses for RT-qPCR data were performed using the one-sided unpaired Student’s t-test with unequal variance. All RT-qPCR data were biologically replicated at least 3 times. All RT-qPCR data are presented as the mean ± s.e.m. as indicated in the figure legends. Statistical analyses for RNA-seq data were performed using the two-sided Mann-Whitney test or the two-sided Wilcoxon signed rank test, as indicated in the figure legend. In Figure 3E, the statistical significance of relative cytosolic RNA expression was calculated using DESeq2 software. In Figure 3H, the statistical significance of the Nuc/Cyt ratio was obtained using the two-sided unpaired Student’s t-test. A *P* < 0.05 was regarded as statistically significant (\**P* < 0.05 \*\**P* < 0.01 \*\*\**P* < 0.001).

## REFERENCES

1. Lander, E.S., Linton, L.M., Birren, B., Nusbaum, C., Zody, M.C., Baldwin, J., Devon, K., Dewar, K., Doyle, M., Fitzhugh, W., et al. (2001). Initial sequencing and analysis of the human genome. Nature 409, 860–921. 10.1038/35057062.

2. Orgel, L.E., and Crick, F.H.C. (1980). Selfish DNA: The ultimate parasite. Nature 284, 604–607. 10.1038/284604a0.

3. Orgel, L.E., Crick, F.H.C., and Sapienza, C. (1980). Selfish DNA. Nature 288, 645–646. 10.1038/288645a0.

4. Kim, Y., Lee, J.H., Park, J.E., Cho, J., Yi, H., and Kim, V.N. (2014). PKR is activated by cellular dsRNAs during mitosis and acts as a mitotic regulator. Genes Dev. 28, 1310– 1322. 10.1101/gad.242644.114.

5. Kim, Y., Park, J., Kim, S., Kim, M.A., Kang, M.G., Kwak, C., Kang, M., Kim, B., Rhee, H.W., and Kim, V.N. (2018). PKR Senses Nuclear and Mitochondrial Signals by Interacting with Endogenous Double-Stranded RNAs. Mol. Cell 71, 1051–1063.e6. 10.1016/j.molcel.2018.07.029.

6. Ahmad, S., Mu, X., Yang, F., Greenwald, E., Park, J.W., Jacob, E., Zhang, C.Z., and Hur, S. (2018). Breaching Self-Tolerance to Alu Duplex RNA Underlies MDA5-Mediated Inflammation. Cell 172, 797–810.e13. 10.1016/j.cell.2017.12.016.

7. Athanasiadis, A., Rich, A., and Maas, S. (2004). Widespread A-to-I RNA editing of Alu-containing mRNAs in the human transcriptome. PLoS Biol. 2, e391. 10.1371/journal.pbio.0020391.

8. Chen, L.L., and Carmichael, G.G. (2009). Altered Nuclear Retention of mRNAs Containing Inverted Repeats in Human Embryonic Stem Cells: Functional Role of a Nuclear Noncoding RNA. Mol. Cell 35, 467–478. 10.1016/j.molcel.2009.06.027.

9. Hu, S. Bin, Xiang, J.F., Li, X., Xu, Y., Xue, W., Huang, M., Wong, C.C., Sagum, C.A., Bedford, M.T., Yang, L., et al. (2015). Protein arginine methyltransferase CARM1 attenuates the paraspecklemediated nuclear retention of mRNAs containing IRAlus. Genes Dev. 29, 630–645. 10.1101/gad.257048.114.

10. Chen, L.L., DeCerbo, J.N., and Carmichael, G.G. (2008). Alu element-mediated gene silencing. EMBO J. 27, 1694–1705. 10.1038/emboj.2008.94.

11. Elbarbary, R.A., Li, W., Tian, B., and Maquat, L.E. (2013). STAU1 binding 3′ UTR IRAlus complements nuclear retention to protect cells from PKR-mediated translational shutdown. Genes Dev. 27, 1495–1510. 10.1101/gad.220962.113.

12. Torres, M., Becquet, D., Blanchard, M.P., Guillen, S., Boyer, B., Moreno, M., Franc, J.L., and François-Bellan, A.M. (2016). Circadian RNA expression elicited by 3’-UTR IRAlu-paraspeckle associated elements. eLife 5. 10.7554/eLife.14837.

13. Schonborn, J., Oberstraß, J., Breyel, E., Tittgen, J., Schumacher, J., and Lukacs, N. (1991). Monoclonal antibodies to double-stranded RNA as probes of RNA structure in crude nucleic acid extracts. Nucleic Acids Res. 19, 2993–3000. 10.1093/nar/19.11.2993.

14. Park, J.E., Yi, H., Kim, Y., Chang, H., and Kim, V.N. (2016). Regulation of Poly(A) Tail and Translation during the Somatic Cell Cycle. Mol. Cell 62, 462–471. 10.1016/j.molcel.2016.04.007.

15. Tian, B., and Manley, J.L. (2016). Alternative polyadenylation of mRNA precursors. Nat. Rev. Mol. Cell Biol. 18, 18–30. 10.1038/nrm.2016.116.

16. Friedman, R.C., Farh, K.K.H., Burge, C.B., and Bartel, D.P. (2009). Most mammalian mRNAs are conserved targets of microRNAs. Genome Res. 19, 92–105. 10.1101/gr.082701.108.

17. Agarwal, V., Bell, G.W., Nam, J.W., and Bartel, D.P. (2015). Predicting effective microRNA target sites in mammalian mRNAs. eLife 4. 10.7554/eLife.05005.

18. Kavakiotis, I., Alexiou, A., Tastsoglou, S., Vlachos, I.S., and Hatzigeorgiou, A.G. (2022). DIANA-miTED: A microRNA tissue expression database. Nucleic Acids Res. 50, D1055– D1061. 10.1093/nar/gkab733.

19. Herrmann, C.J., Schmidt, R., Kanitz, A., Artimo, P., Gruber, A.J., and Zavolan, M. (2020). PolyASite 2.0: A consolidated atlas of polyadenylation sites from 3′ end sequencing. Nucleic Acids Res. 48, D174–D179. 10.1093/nar/gkz918.

20. Gianfrancesco, O., Bubb, V.J., and Quinn, J.P. (2017). SVA retrotransposons as potential modulators of neuropeptide gene expression. Neuropeptides 64, 3–7. 10.1016/j.npep.2016.09.006.

21. Yao, C., Biesinger, J., Wan, J., Weng, L., Xing, Y., Xie, X., and Shi, Y. (2012). Transcriptome-wide analyses of CstF64-RNA interactions in global regulation of mRNA alternative polyadenylation. Proc. Natl. Acad. Sci. U. S. A. 109, 18773–18778. 10.1073/pnas.1211101109.

22. Zhang, S., Zhang, X., Lei, W., Liang, J., Xu, Y., Liu, H., and Ma, S. (2019). Genome-wide profiling reveals alternative polyadenylation of mRNA in human non-small cell lung cancer. J. Transl. Med. 17, 1–11. 10.1186/s12967-019-1986-0.

23. Liberzon, A., Birger, C., Thorvaldsdóttir, H., Ghandi, M., Mesirov, J.P., and Tamayo, P. (2015). The Molecular Signatures Database Hallmark Gene Set Collection. Cell Syst. 1, 417–425. 10.1016/j.cels.2015.12.004.

24. Liu, T., Zhang, L., Joo, D., and Sun, S.C. (2017). NF-κB signaling in inflammation. Signal Transduct. Target. Ther. 2, 1–9. 10.1038/sigtrans.2017.23.

25. Zamanian-Daryoush, M., Mogensen, T.H., DiDonato, J.A., and Williams, B.R.G. (2000). NF-κB Activation by Double-Stranded-RNA-Activated Protein Kinase (PKR) Is Mediated through NF-κB-Inducing Kinase and IκB Kinase. Mol. Cell. Biol. 20, 1278–1290. 10.1128/mcb.20.4.1278-1290.2000.

26. Riley, T., Sontag, E., Chen, P., and Levine, A. (2008). Transcriptional control of human p53-regulated genes. Nat. Rev. Mol. Cell Biol. 9, 402–412. 10.1038/nrm2395.

27. Kulikov, R., Letienne, J., Kaur, M., Grossman, S.R., Arts, J., and Blattner, C. (2010). Mdm2 facilitates the association of p53 with the proteasome. Proc. Natl. Acad. Sci. U. S. A. 107, 10038–10043. 10.1073/pnas.0911716107.

28. Oliner, J.D., Kinzler, K.W., Meltzer, P.S., George, D.L., and Vogelstein, B. (1992). Amplification of a gene encoding a p53-associated protein in human sarcomas. Nature 358, 80–83. 10.1038/358080a0.

29. Hill, D.E., Burrell, M., Oliner, J.D., Sidransky, D., Kinzler, K.W., and Vogelstein, B. (1993). p53 Mutation and MDM2 Amplification in Human Soft Tissue Sarcomas. Cancer Res. 53, 2231–2234.

30. Zhao, K., Yang, Y., Zhang, G., Wang, C., Wang, D., Wu, M., and Mei, Y. (2018). Regulation of the Mdm2–p53 pathway by the ubiquitin E3 ligase MARCH 7. EMBO Rep. 19, 305–319. 10.15252/embr.201744465.

31. Guo, J., Che, X., Wang, X., and Jia, R. (2018). Inhibition of the expression of oncogene SRSF3 by blocking an exonic splicing suppressor with antisense oligonucleotides. RSC Adv. 8, 7159–7163. 10.1039/c7ra11267j.

32. Seo, J.S., Ju, Y.S., Lee, W.C., Shin, J.Y., Lee, J.K., Bleazard, T., Lee, J., Jung, Y.J., Kim, J.O., Shin, J.Y., et al. (2012). The transcriptional landscape and mutational profile of lung adenocarcinoma. Genome Res. 22, 2109–2119. 10.1101/gr.145144.112.

33. Zhang, H., Lee, J.Y., and Tian, B. (2005). Biased alternative polyadenylation in human tissues. Genome Biol. 6, 1–13. 10.1186/gb-2005-6-12-r100.

34. Shepard, P.J., Choi, E.A., Lu, J., Flanagan, L.A., Hertel, K.J., and Shi, Y. (2011). Complex and dynamic landscape of RNA polyadenylation revealed by PAS-Seq. RNA 17, 761–772. 10.1261/rna.2581711.

35. Hilgers, V., Perry, M.W., Hendrix, D., Stark, A., Levine, M., and Haley, B. (2011). Neural-specific elongation of 3′ UTRs during Drosophila development. Proc. Natl. Acad. Sci. U. S. A. 108, 15864–15869. 10.1073/pnas.1112672108.

36. Blair, J.D., Hockemeyer, D., Doudna, J.A., Bateup, H.S., and Floor, S.N. (2017). Widespread Translational Remodeling during Human Neuronal Differentiation. Cell Rep. 21, 2005–2016. 10.1016/j.celrep.2017.10.095.

37. Tsai, Y.L., Coady, T.H., Lu, L., Zheng, D., Alland, I., Tian, B., Shneider, N.A., and Manley, J.L. (2020). ALS/FTD-associated protein FUS induces mitochondrial dysfunction by preferentially sequestering respiratory chain complex mRNAs. Genes Dev. 34, 785–805. 10.1101/gad.335836.119.

38. Xue, Y.C., Ng, C.S., Xiang, P., Liu, H., Zhang, K., Mohamud, Y., and Luo, H. (2020). Dysregulation of RNA-Binding Proteins in Amyotrophic Lateral Sclerosis. Front. Mol. Neurosci. 13, 78. 10.3389/fnmol.2020.00078.

39. Nishimoto, Y., Nakagawa, S., Hirose, T., Okano, H.J., Takao, M., Shibata, S., Suyama, S., Kuwako, K.I., Imai, T., Murayama, S., et al. (2013). The long non-coding RNA nuclear-enriched abundant transcript 1-2 induces paraspeckle formation in the motor neuron during the early phase of amyotrophic lateral sclerosis. Mol. Brain 6, 1–18. 10.1186/1756-6606-6-31.

40. Shelkovnikova, T.A., Kukharsky, M.S., An, H., Dimasi, P., Alexeeva, S., Shabir, O., Heath, P.R., and Buchman, V.L. (2018). Protective paraspeckle hyper-assembly downstream of TDP-43 loss of function in amyotrophic lateral sclerosis. Mol. Neurodegener. 13, 1–17. 10.1186/s13024-018-0263-7.

41. Iridoy, M.O., Zubiri, I., Zelaya, M.V., Martinez, L., Ausín, K., Lachen-Montes, M., Santamaría, E., Fernandez-Irigoyen, J., and Jericó, I. (2019). Neuroanatomical quantitative proteomics reveals common pathogenic biological routes between amyotrophic lateral sclerosis (ALS) and frontotemporal dementia (FTD). Int. J. Mol. Sci. 20, 4. 10.3390/ijms20010004.

42. Smith, E.F., Shaw, P.J., and De Vos, K.J. (2019). The role of mitochondria in amyotrophic lateral sclerosis. Neurosci. Lett. 710, 132933. 10.1016/j.neulet.2017.06.052.

43. Lubelsky, Y., and Ulitsky, I. (2018). Sequences enriched in Alu repeats drive nuclear localization of long RNAs in human cells. Nature 555, 107–111. 10.1038/nature25757.

44. Nakagawa, S., Naganuma, T., Shioi, G., and Hirose, T. (2011). Paraspeckles are subpopulation-specific nuclear bodies that are not essential in mice. J. Cell Biol. 193, 31–39. 10.1083/jcb.201011110.

45. Shalem, O., Sanjana, N.E., Hartenian, E., Shi, X., Scott, D.A., Mikkelsen, T.S., Heckl, D., Ebert, B.L., Root, D.E., Doench, J.G., et al. (2014). Genome-scale CRISPR-Cas9 knockout screening in human cells. Science 343, 84–87. 10.1126/science.1247005.

46. Sanjana, N.E., Shalem, O., and Zhang, F. (2014). Improved vectors and genome-wide libraries for CRISPR screening. Nat. Methods 11, 783–784. 10.1038/nmeth.3047.

47. Harrow, J., Frankish, A., Gonzalez, J.M., Tapanari, E., Diekhans, M., Kokocinski, F., Aken, B.L., Barrell, D., Zadissa, A., Searle, S., et al. (2012). GENCODE: The reference human genome annotation for the ENCODE project. Genome Res. 22, 1760–1774. 10.1101/gr.135350.111.

48. Kim, D., Paggi, J.M., Park, C., Bennett, C., and Salzberg, S.L. (2019). Graph-based genome alignment and genotyping with HISAT2 and HISAT-genotype. Nat. Biotechnol. 37, 907–915. 10.1038/s41587-019-0201-4.

49. Li, H., Handsaker, B., Wysoker, A., Fennell, T., Ruan, J., Homer, N., Marth, G., Abecasis, G., and Durbin, R. (2009). The Sequence Alignment/Map format and SAMtools. Bioinformatics 25, 2078–2079. 10.1093/bioinformatics/btp352.

50. Liao, Y., Smyth, G.K., and Shi, W. (2019). The R package Rsubread is easier, faster, cheaper and better for alignment and quantification of RNA sequencing reads. Nucleic Acids Res. 47, e47–e47. 10.1093/nar/gkz114.

51. Love, M.I., Huber, W., and Anders, S. (2014). Moderated estimation of fold change and dispersion for RNA-seq data with DESeq2. Genome Biol. 15, 1–21. 10.1186/s13059-014-0550-8.

52. Wu, T., Hu, E., Xu, S., Chen, M., Guo, P., Dai, Z., Feng, T., Zhou, L., Tang, W., Zhan, L., et al. (2021). clusterProfiler 4.0: A universal enrichment tool for interpreting omics data. Innovation 2, 100141. 10.1016/j.xinn.2021.100141.

53. Yu, G., Wang, L.G., Yan, G.R., and He, Q.Y. (2015). DOSE: An R/Bioconductor package for disease ontology semantic and enrichment analysis. Bioinformatics 31, 608–609. 10.1093/bioinformatics/btu684.

54. Gruber, A.R., Lorenz, R., Bernhart, S.H., Neuböck, R., and Hofacker, I.L. (2008). The Vienna RNA websuite. Nucleic Acids Res. 36, W70–W74. 10.1093/nar/gkn188.

55. McGeary, S.E., Lin, K.S., Shi, C.Y., Pham, T.M., Bisaria, N., Kelley, G.M., and Bartel, D.P. (2019). The biochemical basis of microRNA targeting efficacy. Science 366. 10.1126/science.aav1741.

56. Lim, J., Ha, M., Chang, H., Kwon, S.C., Simanshu, D.K., Patel, D.J., and Kim, V.N. (2014). Uridylation by TUT4 and TUT7 marks mRNA for degradation. Cell 159, 1365–1376. 10.1016/j.cell.2014.10.055.

57. Madeira, F., Pearce, M., Tivey, A.R.N., Basutkar, P., Lee, J., Edbali, O., Madhusoodanan, N., Kolesnikov, A., and Lopez, R. (2022). Search and sequence analysis tools services from EMBL-EBI in 2022. Nucleic Acids Res. 50, W276–W279. 10.1093/nar/gkac240.

