## Supplemental Information for "Alternative Polyadenylation Determines the Functional Landscape of Inverted Alu Repeats"

**SUPPLEMENTARY FIGURE LEGENDS**

**Figure S1, related to Figure 1. Characterization of functional 3′ UTR IRAlus genes.** (A) Successful subcellular fractionation (Cytoplasm: TUBB, GAPDH, and Rab5; Nucleus: Lamin A/C) confirmed by western blotting in HeLa cells. (B and C) The Nuc/Cyt ratio (B) or the translational efficiency (C) of mRNAs with indicated J2 enrichment score bins in HeLa cells. Correlation *r* and its *P* was calculated using the Spearman correlation. (D) Relative abundance of miRNAs in HeLa cells based on DIANA-miTED. miRNAs with an abundance of more than 1% of total miRNA abundance (*n* = 22) were indicated in red. (E and F) Probability of conserved targeting (Aggregate P_CT_; E) and targeting efficacy (CWCS; F) of selected miRNAs from (D) on mRNAs with indicated J2 enrichment scores. The *P* was calculated using the two-sided Mann-Whitney test. (G) Cumulative distribution of the 3′ UTR length of mRNAs with high or low J2 enrichment scores. (H) The half-life of mRNAs with high or low J2 enrichment scores. The *P* was calculated using the two-sided Mann-Whitney test. Boxplots show the 25th, 50th, and 75th percentiles. Whiskers represent a 1.5× interquartile range.

**Figure S2, related to Figure 2. Characteristics of noIRAlus genes with high J2 enrichment scores.** (A) Representative example of noIRAlus genes with dsRNA structure consisting of LINE (*NBPF3*). (B) Gene structure of SVA. SVAs are composed of flanked target site duplications (TSDs), CCCTCT hexamer repeats, an inverted Alu-like sequence, a GC-rich variable number tandem repeat (VNTR), a SINE-R domain, and poly(A) tail. (C) Representative example of interaction between Alu and SVA (*TRIM56*) and IGV snapshot of J2 fCLIP-seq reads mapped to the *TRIM56* 3′ UTR. RNAfold prediction of RNA secondary structure is shown. (D) The Nuc/Cyt ratio of *TRIM56* mRNA analyzed by RT-qPCR (*n* = 6). The *P* was calculated using the one-sided Student’s t-test with unequal variance. (E) Cumulative distribution of the Nuc/Cyt ratios of mRNAs with the indicated 3′ UTR IRAlus. The *P* was calculated using the two-sided Mann-Whitney test. Boxplots show the 25th, 50th, and 75th percentiles. Whiskers represent a 1.5× interquartile range. **P* < 0.05, ***P* < 0.01, ****P* < 0.001.

**Figure S3, related to Figure 3. The downstream effect of 3′ UTR lengthening.** (A) Knockdown efficiency analyzed by RT-qPCR in A549 cells transfected with a mixture of siCSTF2 and siCSTF2T for 4 days (*n* = 3). (B) The J2 enrichment scores in A549 cells transfected with siLuc or a mixture of siCSTF2 and siCSTF2T for 4 days. For the threshold, the median J2 enrichment score of IRAlus genes in A549 cells transfected with siLuc was used. (C) Activation of PKR/eIF2α pathway in A549 cells transfected with a mixture of siCSTF2 and siCSTF2T for 4 days analyzed by western blotting. (D) Relative expression of DEGs listed in HALLMARK_P53_PATHWAY gene set in A549 cells transfected with a mixture of siCSTF2 and siCSTF2T for 4 days (*n* = 3).

**Figure S4, related to Figure 5. Regulation of MDM2 expression by IRAlus.** (A) Sanger sequencing result of PCR amplicons of the 3′ UTR of *MDM2* gene region from gDNAs of A549 WT or A549 ΔIRR cells. Sequences of sgRNAs were indicated in red or blue. (B) PCR amplicon from single cell clones of A549 ΔIRR cells analyzed on a 1% TAE agarose gel. The expected migration of PCR amplicons is shown on the right with the expected size indicated. (C) PCR amplicon from multiplex PCR analysis performed using cDNAs prepared from A549 WT or A549 ΔIRR cells transfected with a mixture of siCSTF2 and siCSTF2T for 4 days analyzed on a 3% TBE agarose gel. The expected migration of PCR amplicons is shown on the right with the expected size indicated. (D) Nuc/Cyt ratio change of *MDM2* mRNA in A549 WT or A549 ΔIRR cells after transfection with a mixture of siCSTF2 and siCSTF2T for 4 days (*n* = 3). Data are represented as mean ± s.e.m. (E) CCK-8 cell viability test in A549 WT or A549 ΔIRR cells transfected with a mixture of siCSTF2 and siCSTF2T for 4 days. (F) PCR amplicon from multiplex PCR analysis performed using cDNAs prepared from A549 WT cells transfected with an ASO targeting the CSTF2 binding sites of *MDM2* mRNA for 2 days. (G) Relative RNA expression of different genomic positions of *MDM2* (CDS, cUTR, and aUTR) analyzed by RT-qPCR in A549 WT or A549 ΔIRR cells transfected with an ASO targeting the CSTF2 binding sites of *MDM2* mRNA (*n* =3). Data are represented as mean ± s.e.m. The *P* was calculated using the one-sided Student’s t-test with unequal variance. n.s. not significant, **P* < 0.05, ***P* < 0.01, ****P* < 0.001.

**Figure S5, related to Figure 6. Regulation of the p53 pathway in lung adenocarcinomas.** (A) Relative *CSTF2* expression in human lung adenocarcinomas when compared to those of their adjacent matched-normal tissues (*n* = 69). The *P* was calculated using the two-sided Mann-Whitney test. (B) Result of GSEA for the relative gene expression in human lung adenocarcinomas with high CSTF2 expression on the HALLMARK_P53_PATHWAY gene set. (C) Heatmap of APA level of *MDM2* (*MDM2* aUTR/CDS) and relative expression of genes previously identified as directly induced or repressed by the p53 pathway in human lung adenocarcinoma tissues (*n* = 69).

**Figure S6, related to Figure 7. Examples of NPC-specific gene silencing by IRAlus.** (A) IGV snapshot of sequencing read accumulation at *NEAT1* locus after subcellular fractionation in NPCs. (B) IGV snapshot of J2 fCLIP-seq read accumulation at the *RPL15* locus in HeLa and NPCs. Alu elements are indicated with arrows. (C and D) Over-representation analysis of genes involved in the KEGG ALS pathway in the Gene Ontology Biological Process (BP; C) or Cellular Component (CC; D) using clusterProfiler. (E-G) IGV snapshot of J2 fCLIP-seq reads mapped to *NUDFS5* (E), *COX6B1* (F), and *UQCFB* (G) gene loci in HeLa and NPCs. These genes were shown as they were involved in the KEGG ALS pathway with NPC-specific gene silencing.

**Figure S7, related to Figure 1. Identification of 3′ UTR IRAlus genes.** (A) Length distribution of Alu elements located in the human genome or just in 3′ UTRs. (B) *P* of Nuc/Cyt ratio comparison between mRNAs of genes with or without 3′ UTR IRAlus after varying the maximum distance between two closest inverted Alu elements. (C) Distribution (left) and cumulative density (right) of the length of 3′ UTR (*n* = 17,515) and distance between two inverted Alu elements located in 3′ UTRs (*n* = 5,551). For distribution, the most frequent 3′ UTR length (315 nt) and distance between two inverted Alu elements (508 nt) are marked. The median 3′ UTR length (1,039 nt) and distance between two inverted Alu elements (1,557 nt) are indicated. (D) Overview and the result of the 3′ UTR IRAlus gene identification.

**Table S1.** Sequences of siRNAs and ASOs

| **Gene** | **Sense (5′-3′)** | **Antisense (5′-3′)** |
| --- | --- | --- |
| siLuc | CUU ACG CUG AGU ACU UCG A | UCG AAG UAC UCA GCG UAA G |
| siCSTF2_1 | CGU UCU CUA CGU UCU GUG U | ACA CAG AAC GUA GAG AAC G |
| siCSTF2_2 | GUC AUG CAG GGA ACA GGA A | UUC CUG UUC CCU GCA UGA C |
| siCSTF2T_1 | GAU AAC UCG CGU UUA CAU A | UAU GUA AAC GCG AGU UAU C |
| siCSTF2T_2 | CUG GUU UAA UGU UGC UUC A | UGA AGC AAC AUU AAA CCA G |
| siNEAT1 | [Phosphate]GUG AGA AGU UGC UUA GAA ACU UUdC dC | GGA AAG UUU CUA AGC AAC UUC UCA CUU |
| siMDM2 | GGA AGA AAC CCA AGA CAA A | UUU GUC UUG GGU UUC UUC C |
| non-target ASO | mA*mC*mU*mC*mU*mA*mU*mC*mU*mG*mC*mA*mC*mG*mC*mU*mG*mA*mC*mU* |  |
| ASO #1 | mA*mA*mU*mU*mG*mC*mA*mU*mU*mC*mU*mU*mG*mA*mA*mA*mC*mA*mA*mU*mU*mC*mU*mU*mA* |  |
| ASO #2 | mA*mA*mC*mA*mA*mC*mU*mC*mA*mU*mA*mA*mA*mA*mA*mU*mU*mU*mA*mA*mG*mA*mU*mC*mA* |  |

*Lowercase “m” designates the 2′-OMe modification. All ASOs have a phosphorothioate-modified backbone at every position.

**Table S2.** Sequences of primers for RT-qPCR and multiplex RT-PCR

| **Gene** | **Forward Primer (5′-3′)** | **Reverse Primer (5′-3′)** |
| --- | --- | --- |
| GAPDH | CTC CTC CAC CTT TGA CGC TG | TCC TCT TGT GCT CTT GCT GG |
| ACTB | CCT GTA CGC CAA CAC AGT GC | ATA CTC CTG CTT GCT GAT CC |
| NEAT1 | GGA GGG CCG GGA GGG CTA AT | CGG TCA GCC CCG TCG AGC TA |
| GATD1 | ACT TCG TGA AGG ATT CGG GC | GAC AGT GGA GCT GGC ATT CT |
| METTL7A | AGC GGG AGC TCT TCA GTA AC | TCT CAA AGT TGG GGT TGG GG |
| MAVS | CTG CTT CGA GGA TCT TGC CA | CAA AGG TGC CCT CGG ACT TA |
| MDM4 | GAG GAG TGG GAT GTA GCT GG | CCC ACT TCA ATC ACC TGA TTT GTC |
| RPL13 | TTC GGT ACC ACA CGA AGG TG | GGA CTC CGT GGA CTT GTT CC |
| RPS19 | AAC TGG TTC TAC ACG CGA GC | TCT TGG TCA TGG AGC CAA CC |
| RPL27A | AGC TTC TGC CCA ACT GTC AA | ATC AAT GAT GGG AGC AGC CC |
| RPL37 | GGA ACT GGT CGA ATG AGG CA | CAG CTG CCC TCT TGG GTT TA |
| DSTN | ACC AGA AGA TCT CAA TCG GGC | GGC ATC CTT CAA AGG CTA CAA |
| TRIM56 | CAC CTT CTT CGG CTG TCC TT | CAG TCT TCG GCT CAT CCT CC |
| KRBA2 | TCA TGT CAC TGG GCC AAA GG | TGC AAG GAA ACG TCG AAA GC |
| ZNF791 | TGT GAC GAA GAA GAC TGC CG | ATG AGA GAC GCA TGA AGG CT |
| PDP2 | GTG GCA AAT GCT GGT GAC TG | ACG TGT AAG GGG CAG ACA AG |
| HAUS3 | TGG TTG TTT GAG GGC GTT GA | ATC CAA TGC CGC CCC TTC TA |
| CSTF2 | GCA AGG AAC CCT ACA GCA CT | CTC GAG GGG TTG GGA CAT TC |
| CSTF2T | TCT GTT GTC AGT TTC CGG CT | GAG GTT CCG CAT GGC ACT AA |
| MDM2_CDS | GCC CTT CGT GAG AAT TGG CT | AAG CCC TCT TCA GCT TGT GTT |
| MDM2_cUTR | AGG AGT ATC GGT AGC ATA AAT GTG A | TAA AAA TAG CTG TCA CTG CCT CCA |
| MDM2_aUTR | GCA GAT GTA AGC TTG AGC CC | ACT AAC AGT CCT AAC CCC TGC |
| P21 | AGG TGG ACC TGG AGA CTC TCA G | TCC TCT TGG AGA AGA TCA GCC G |
| PUMA | CGA CCT CAA CGC ACA GTA CGA | AGG CAC CTA ATT GGG CTC CAT |
| BAX | TGG AGC TGC AGA GGA TGA TTG | GAA GTT GCC GTC AGA AAA CAT G |
| Multiplex RT-PCR | #1 AGG AGT ATC GGT AGC ATA AAT GTG A | #2 TAA AAA TAG CTG TCA CTG CCT CCA  #3 ATT TCT AAA GTG GGC AGG CA |

**Table S3.** Sequences of primers for *in vitro* transcription

| **Primer** | **Forward Primer (5′-3′)** | **Reverse Primer (5′-3′)** |
| --- | --- | --- |
| ZNF587 | AGG CTC ACC CCT GTG AAA TG | TAA TAC GAC TCA CTA TAG GGC ACG CTG TGC TTA TTG CCA AA |
| MCM4 | CAA AGG CTG GGA TCA TCT GT | TAA TAC GAC TCA CTA TAG GGC AGC TAC TCG GGA GGT TGA G |
| RPL13 | TTG TGG GGA GGT TAC AGA GG | TAA TAC GAC TCA CTA TAG GGC GGC CCA CCT TAA CTT TAC A |
| RPL15 | GTA AGC CAA GAT GGG TGC AT | TAA TAC GAC TCA CTA TAG GGG GGA GTG GAC AGA GTG GGT A |
| NEAT1 | GGA GGG CCG GGA GGG CTA AT | TAA TAC GAC TCA CTA TAG GGG CTA AGG GGC AGC GAA GGA TGC |
| GAPDH | CAA ATT CCA TGG CAC CGT CAA | TAA TAC GAC TCA CTA TAG GGT TAC TCC TTG GAG GCC ATG TG |
| GATD1 | CCC AGT CCT TCC TCC ACT GT | TAA TAC GAC TCA CTA TAG GGC CTT CAC GAA GTC CTC CAC C |
| PDP2 | CAG TGT TGC GGT TTG AGA GC | TAA TAC GAC TCA CTA TAG GGC CTC GGT ATT GAA GCC CCT C |
| DSTN | TCG TAA ATG CTC CAC ACC AGA A | TAA TAC GAC TCA CTA TAG GGA TTG CAT CCT TGG AGC TTG C |
| MDM2 | ATG GTG AGG AGC AGG CAA AT | TAA TAC GAC TCA CTA TAG GGT CTA CAT ACT GGG CAG GGC T |

**Table S4.** Sequences of sgRNAs for CRISPR-Cas9

| **Gene** | **Forward Primer (5′-3′)** | **Reverse Primer (5′-3′)** |
| --- | --- | --- |
| sgMDM2_1 | CAC CGG AGG TCA AGG TCA AGA CAC G | AAA CCG TGT CTT GAC CTT GAC CTC C |
| sgMDM2_2 | CAC CGG CTA TGG CAA GAA GAT ACA A | AAA CTT GTA TCT TCT TGC CAT AGC C |
| sgMDM2_3 | CAC CGA AGT CTA GGT AAA TAT ACC A | AAA CTG GTA TAT TTA CCT AGA CTT C |
| sgMDM2_4 | CAC CGT TAG TCA TAG GAC AGA GGA T | AAA CAT CCT CTG TCC TAT GAC TAA C |

**Table S5.** Sequences of primers for gDNA amplification

| **Primer** | **Forward Primer (5′-3′)** | **Reverse Primer (5′-3′)** |
| --- | --- | --- |
| MDM2 gDNA | ACC CCA AAT CCA GCT TAG GTA GCC | GGG AGA GGC CAT GAC TAC CTC TGA A |
